# A quantitative interactome map reveals affinity tailored hijacking of human cellular functions by resident and pathogenic bacteria

**DOI:** 10.64898/2026.10.02.755967

**Authors:** Jaime Fernandez-Macgregor, Laura Ilari, Fabien Durbesson, Stéphane Audebert, Luc Camoin, Bastien Morlet, Virginie Millet, Jean Michel Dela Vedova-Costa, Charlene Gaedke, Vincent Kourilsky, Lou Bergogne, Eric Martinez, Achille Broggi, Jean-Paul Borg, Søren Østergaard, Gilles Trave, Elodie Monsellier, Christine Brun, Andreas Zanzoni, Renaud Vincentelli

## Abstract

Bacteria employ molecular mimicry to hijack host cellular functions for their own benefit. Despite the wealth of data mapping human-bacteria interactions, a comprehensive view of how these networks are fine-tuned remains elusive. In particular, there have been no large-scale screenings measuring human-bacterial protein-protein binding affinities to explore their role in functional and pathological outcomes. Here, we report the largest quantitative interactome to date between the family of human PDZ domains and C-terminal PDZ-binding motifs (PBMs) in effectors from pathogenic and resident gut bacteria. We show that bacteria tailor effector sequences to bind PDZ domains with similar or higher binding affinities than human PBMs to hijack their associated cellular functions. For a subset of the dataset, we validate PDZ-PBM interactions with the corresponding full-length proteins and show that for pathogenic *E. coli* effector NleA, its C-terminal PBM is necessary for interactions with proteins involved in intracellular transport and membrane organization, probably impacting the host glycocalyx to favor infection. Finally, we show that motifs overlapping with effector PBMs target interaction modules other than PDZ domains, adding an additional layer of complexity to the host-microbe molecular dialogue.

## Introduction

Bacteria, viruses, fungi and parasites have evolved to establish a molecular dialogue with their hosts, that is often mediated by effector proteins. These proteins can be expressed in or translocated into host cells, where they interact with host proteins to hijack their associated cellular functions^1–5^. Perturbation of host protein-protein interactions (PPIs) has been amply documented for both plants^6^ and humans^7–13^, and often involves molecular mimicry by microorganismal effectors, that imitate host short linear motifs (SLiMs)^5,14,15^, globular domains and full proteins^16,17^.

Among these microorganisms, Gram-negative bacteria of the Pseudomonadota phylum are associated with infectious diseases in mammals and plants^18^ and occur naturally in the gut microbiota, representing 5-15% of all bacteria therein^19^. The higher end of this relative abundance spectrum is associated with dysbiosis, inflammation, metabolic disorders and cancer^19^. Recently, we showed that gut resident Pseudomonadota bacteria encode complete and functional type III secretion systems (T3SS) with which they can translocate effectors into human cells, some of which interact with human PDZ domains via C-terminal PDZ-binding motifs (PBMs)^20^. SLiM mimicry is also prevalent among pathogenic Pseudomonadota virulence factors secreted via T3SS, T4SS and T6SS^17,21–24^ and plays a crucial role in the subversion of host defenses^5,14,16^. In some cases, effector motifs are implicated in cancer^25^ and display tailored binding affinities and specificities for human protein targets to disrupt signaling pathways^26^.

Effectors secreted via T3SS by pathogenic strains of *E. coli*^27–31^, *Shigella*^32^ and *Salmonella*^33^ are enriched in C-terminal PDZ-binding motifs^32^ (PBMs) and target human PDZ domains (Figure 1a) . Human PDZ-mediated activities of *E. coli* effectors include cell surface projection (Map, EspI)^27^, localization to the Golgi (EspI/NleA)^28^, disruption of tight-junctions (EspG1/G2, NleA, EspF and Map)^31^ and NF-κB pathway activity (NleH)^29^. *Shigella* effector OspE promotes protein kinase C activation via PDLIM proteins in a PDZ dependent manner^32^. Further, *in celllula* interactions of essential virulence effectors such as NleA from enteropathogenic *E. coli* (EPEC) strains have been shown to be mediated by its C-terminal PBM, the truncation of which attenuates virulence^28^. During infection, EPEC forms attaching and effacing lesions on human intestinal mucosa via secreted effectors and causes a perturbation of the epithelial barrier^28,34,35^. The involvement of PDZ-containing proteins in cellular junctions, cellular adhesion, cell polarity, transmembrane transport, signaling and actin related processes^36,37^ makes them relevant targets.

**Figure 1.**
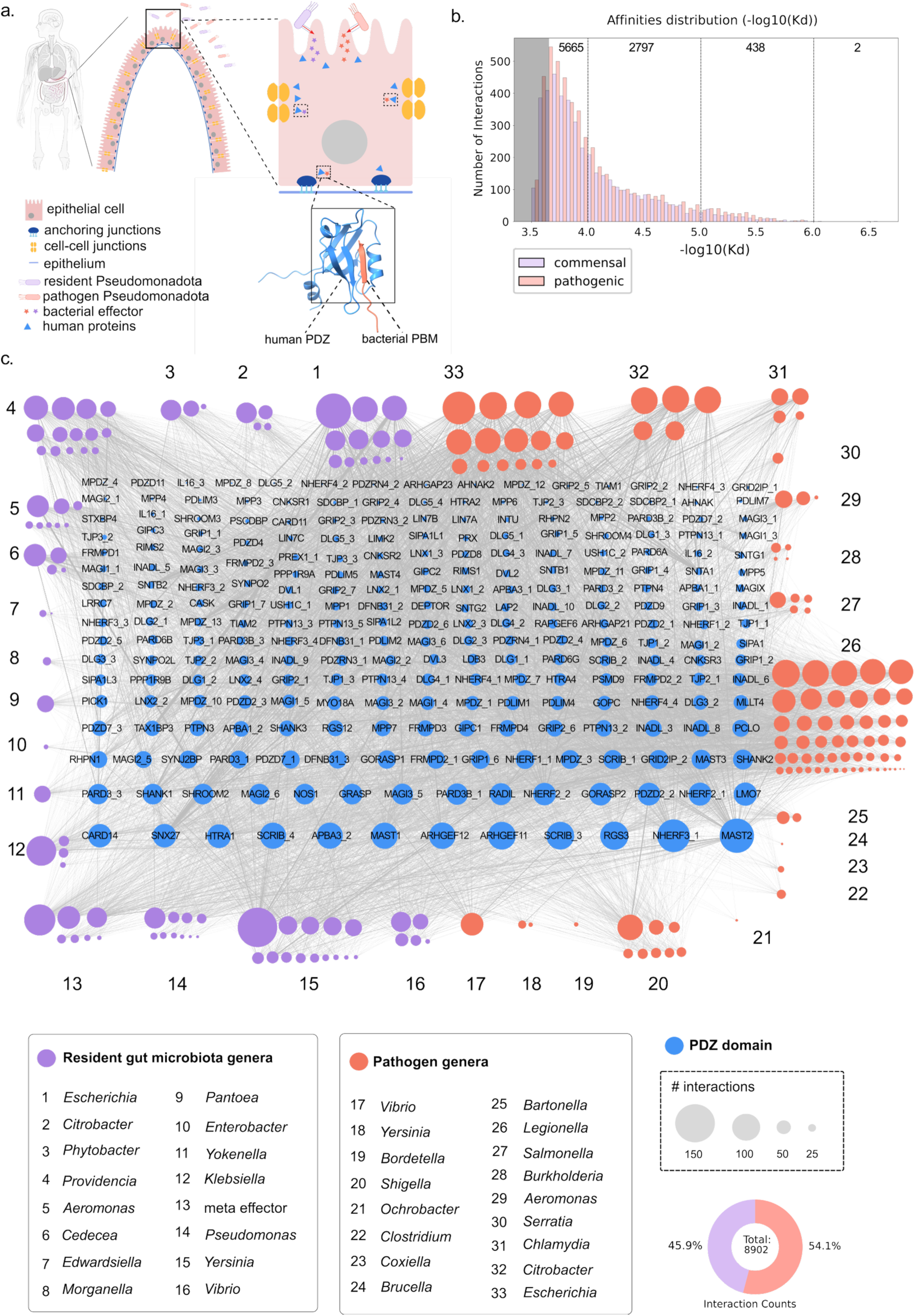
PDZ-PBM interaction network. **a,** Illustration of effector secretion into intestinal epithelial cells, with an example PDZ-PBM interface showing characteristic β-strand augmentation upon binding **b,** Distribution of binding affinities (-log(Kd)) for C-terminal PBMs in effectors from resident gut microbiota strains (purple) and known pathogens (salmon). Number of significant interactions for each affinity interval are indicated inside plot area at the top. The grey area on the left indicates the interval for which variability between measurements is higher and recovery lower, as also observed in^41^ **c,** PDZ-PBM interaction network showing all PDZ domains (blue nodes) in the central grid, arranged according to number of interactions (node degree) and PBMs grouped and numbered by genus of origin surounding the central grid. Node sizes correspond to number of interactions as shown in the legend on the bottom right. Resident gut genera are shown as purple nodes and pathogen ones as orange nodes. Meta effector (c, number 13) are effectors from meta genome assembled genomes^44^.

Despite growing evidence of the importance of effector SLiMs in bacterial infection and their potential to outcompete human proteins for binding partners, there have been no large scale quantitative interactome maps between human domains and bacterial SLiMs, although recent high-throughput methods show promise^38,39^.

Here, we report the largest quantitative interactome to date for bacterial SLiMs and human globular domains, covering a space of ∼50,000 possible interactions. We report binding affinities for ∼9,000 (∼18%) interactions between the full family of human PDZ domains (PDZome, comprising 266 domains distributed across 152 proteins) and ∼200 C-terminal PBM (10 amino acids long) of secreted effectors from 38 and 34 strains of resident microbiota and pathogenic bacteria, respectively (Fig.1).

### A quantitative interactome between human PDZ domains and bacterial C-terminal PDZ-binding motifs

First, we identified all SLiMs matching entries in the ELM database^40^ present in effector proteins secreted via T3SS and T4SS and found C-terminal PBMs matching ELM classes (LIG_PDZ_Class_1, 2, 3 and Wminus1_1, see Methods) in 849/4183 of them (∼20%) (Figure S1). We selected ∼200 ten amino acids long bacterial PBMs (see Methods), including well characterized ones from known pathogens^27,32^.

We produced and purified 253/266 human PDZ domains and quantified binding against our library of bacterial peptides using the high-throughput fragmentomic Holdup assay, a quantitative *in vitro* interaction assay we previously developed^41,42^. Briefly, biotinylated PBMs are bound on streptavidin coated beads, incubated until equilibrium with purified PDZ domains, then the unbound fraction is quantified and converted to a dissociation constant (Kd) (see Methods).

Out of ∼50,000 binary PDZ-PBM interactions tested, we report binding affinities above our sensitivity threshold for ∼9000 of them (18% of all tested pairs) with dissociation constants (Kd) in the ∼0.25-330µM range (Fig. 1,b). To our knowledge, this is the largest quantitative interactome to date between human domains and bacterial SLiMs (Fig. 1, c).

To control the quality of positive hits, we systematically included 24 positive controls from the Profaff database^41^, the largest quantitative interactome in literature, reporting ∼18,000 dissociation constants (Kd) between human PDZ domains and human and viral PBMs. We measured ∼3,300 interactions that are reported there, showing a good correlation (Pearson’s R=0.88), with a few mismatches attributable to a limited number of PDZs and PBMs (Figure S2). The overall sensitivity, specificity and precision of the assay are at least ∼70% for the lower end of the affinity range and increase up to ∼95% for higher affinities (Figure S3). Hence, the selected threshold represents a good compromise for discovery and is amply backed up by our positive controls.

As an additional validation, we downloaded data from the IntAct database^43^ for all experimentally confirmed PPIs with interface annotations between human proteins and bacterial effectors corresponding to all tested PBMs. Out of 17 interactions for which interface data is available in IntAct, 15 report either a PDZ or a PBM in the interface (6 PBM only, 4 PDZ only and 5 both), 13 (∼87%) for which we quantified the corresponding PDZ-PBM interaction (Figure S4).

Interactions were quantified for PBMs from all 72 tested strains, with the total number per strain varying from 3 (*Edwardsiella tarda)* to 2079 (*Legionella pneumophila)* (Figure 1c and S5). For each strain, some PBMs were more promiscuous than others, with the lowest binder having 2 interactions and the highest one 145 (Figure S5), with the least promiscuous binders often harboring an Alanine at the C-terminal. PBMs from *Shigella flexneri, Citrobacter rodentium*, *Vibrio parahaemolyticus,* resident *E. coli (*MS 155-1, 69-1, 196-1), pathogenic *E.coli* and *Escherichia fergusonii* effectors are the most promiscuous on average, with >50 interactions per PBM (Figure S6).

Human PDZ domains have as few as 1 (pro-interleukin 16 (IL16) third PDZ domain) and as many as 127 (microtubule associated serine threonine kinase 2 (MAST2)) significant interactions (Figure 1c). MAST2 is also the most promiscuous binder with human proteome PBMs^41^, and more broadly, PDZ domains that are network hubs in the human PDZ-PBM interactome^41^ are also hubs in the human-bacterial interactome, as shown by the linear correlation of normalized node degrees for both datasets (Figure S7). Additionally, PDZs show wide strain promiscuities, with some targeted by >20 strains and others by as few as 1 (Figure S8). Overall, our data indicate PDZ domains interact with resident and pathogenic bacterial PBMs and may be relevant to the host-microbe molecular dialogue.

### Bacteria tailor effector sequences to outcompete human PDZ-PBM interactions and hijack specific cellular functions

Assessing if the quantified PDZ-PBM interactions are likely to be biologically relevant requires a multi-step approach. First, it is necessary to determine what the relationship between binding affinities and biological functions is within the human PDZ domain family, to then broaden the scope to include bacterial PBMs. Then, by comparing the strongest binders for PDZ domains with specific functions, we can propose human PDZ-PBM mediated interactions likely to be perturbed by bacterial effectors via affinity driven interaction displacement.

To this end, we clustered all tested bacterial PBMs with the human PBMs we previously published interactions for^41^, based on their interaction similarity profiles for human PDZ domains. These were calculated using affinity vectors for each PBM from the affinity weighted adjacency matrix of our measured interactome, for all high-confidence interactions (pKd>4, Figure 1b and S2). Then, we performed functional enrichment of proteins in the resulting clusters (Figure 2a-c) and calculated binding energy differences for each bacterial and human PBM compared with the highest affinity human PBM for each PDZ domain (Figures 2d and S9) (see Methods).

**Figure 2.**
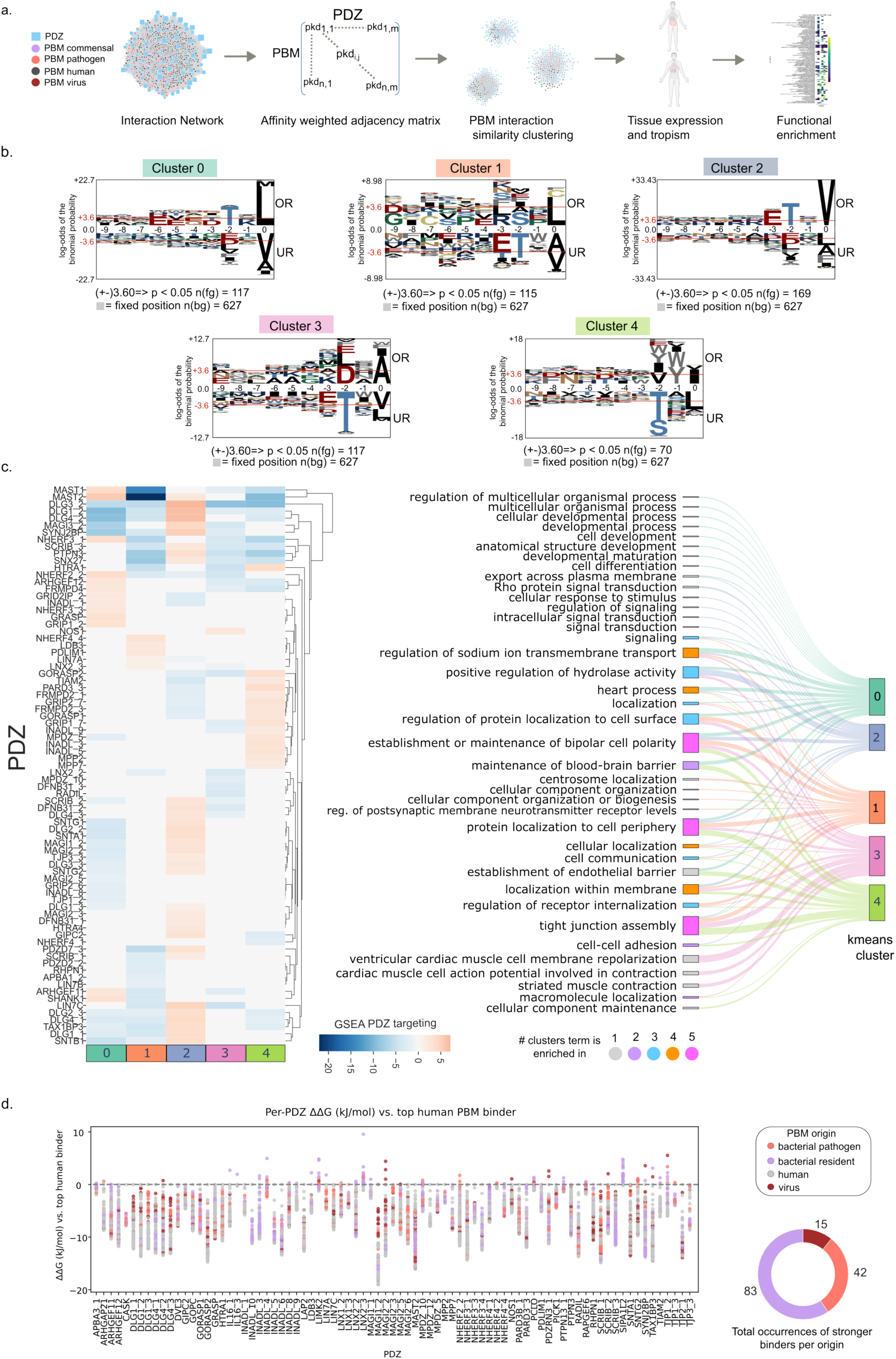
Functional enrichment of interaction clusters and ΔΔG comparisons. **a,** diagram illustrating PBM clustering and functional enrichment workflow. **b,** position specific scoring matrix logos for 10 amino acid PBMs in each interaction similarity Kmeans cluster, for interactions with pkd >4, showing over and under represented (OR and UR, respectively) amino acids at each position from the C-terminus (right side of each logo). Red horizontal lines shown at +3.6 and -3.6 on the Y-axes correspond to *P-value <0.05* (done with pLogo^53^) **c,** (left) GSEA heatmap with normalized enrichment scores for preferential targeting of PDZ domains by members of each cluster (calculated based on ranked affinities) and (right) alluvial plot showing significantly enriched (Benjamini-Hochberg *FDR* < 10^-2^) Gene Ontology Biological Process terms (GO:BP) for interaction similarity clusters for human proteins expressed in GI tract tissues and effectors from bacteria with GI tissue tropism. Nodes for GO:BP terms are colored according to the total number of clusters in which the term is enriched. Semantic similarity clustering of enrichment terms was performed with Revigo^54^ (threshold 0.7, *Homo sapiens*). **d,** (left) scatter plot of differences in binding energies (ΔΔG in kJ/mol) for PBMs from bacteria with gut tropism and human proteins expressed in gut tissues, with respect to the human PBM with the highest affinity for each PDZ domain and (right) donut plot showing number of instances where a PBM of non-human origin has a higher affinity than the top human PBM binder for all PDZ domains.

Clustering revealed five groups with distinct amino acid compositions (Figure 2b), with PBMs from multiple pathogen and resident gut strains present in each one (Figure S10). Gene set enrichment analysis shows preferential targeting of specific PDZ domains by members of each cluster (Figure 2c, heatmap) and functional enrichment analysis reveals shared and exclusive functions enriched in each cluster (Figure 2c, alluvial plot). For certain GO Biological Process terms enriched in more than one cluster, different proteins drive the enrichment, implying common functions could be perturbed via different targets.

For instance, while establishment and maintenance of cell polarity is enriched in every cluster (Figure 2c, alluvial plot), we noticed that PDZ domains in proteins from baso-lateral (SCRIB-DLG1-LGL) and apical (PAR3/PAR6/aPKC and CRUMBS/PALS1/PATJ) polarity complexes^45^, which are crucial for establishment and maintenance of cell-cell junctions^46^, are targeted by PBMs in clusters 2 and 4, respectively (Figure 2c, heatmap). Of note, all EPEC NleA PBMs have affinities in the ∼2-5µM range for the 3^rd^ PDZ of Disks large homolog 1 (DLG1), similar to their strongest human PBM binder from Cysteinyl leukotriene receptor 2 (CYSLTR2), that shares the last four C-terminal amino acids with them (`ETRV`) and is strongly expressed in the gut^47^. For Protein scribble homolog (SCRIB), PBMs from all nine NleA homologs and from effectors from resident gut strains *Providencia stuartii* (ATCC 25827), *Escherichia coli* (MS 198-1 and MS 69-1) and *Yersinia enterocolitica* (subsp. palearctica Y11) have a total of 33 interactions with Kd<18 µM with SCRIB domains 1, 2, 3 and 4, potentially impacting its role in cell polarity processes as well as its recruitment to cell membranes and deregulation of its tumor suppressing activities, both of which depend on its PDZ-PBM interactions^45,48^. Similarly, of three Partitioning defective 3 homolog (PARD3) PDZ domains we tested, the third one is also present in the Profaff database^41^. Its top six binders (Kd ∼2-5µM), all in cluster 4, include Transient receptor potential cation channel subfamily V member 6 (TRPV6) which is critical for calcium uptake in epithelial tissues^49^, along with PBMs of effectors from resident *Providencia rettgeri*, *E. fergusonii*, *Cedecea davisae* and *Phytobacter massiliensis* and with slightly lower affinities (Kd ∼6-10 µM) EHEC effector EspX2 and a *Legionella* Dot/Icm T4SS effector. Finally, enrichment in cluster 4 for protein associated to tight junctions (PATJ) PDZ domains is mainly driven by resident gut PBMs, which display higher median affinities for them than PBMs from pathogens (Figure S12). Loss of cell polarity can lead to erosion of the intestinal epithelial layer, a phenomenon observed during infection by pathogenic bacteria and in gut dysbiosis. Targeting of SCRIB, DLG1, PARD3 and PATJ by resident and pathogen effectors alike, with affinities similar to or higher than their strongest human binders, provides likely molecular targets behind perturbation of host cell polarity and points to specific bacterial species that may be associated with disease phenotypes.

Next, we noticed Clusters 0 and 2 contain PBMs from several human solute carriers (SLCs), plasma membrane calcium transporting ATPases and the cystic fibrosis transmembrane regulator (CFTR), a cAMP and cGMP-regulated chloride (Cl^-^) and bicarbonate (HCO3^-^) channel located at apical surfaces of epithelial cells where it participates in ion transport across the cell membrane^50^. Thus, effectors present in this cluster are likely to perturb cellular fluid and electrolyte balance. These human PBMs jointly have 33 interactions with Kd<3µM with 15 PDZ domains, 5 of which are targeted by members of both clusters, and 3 and 7 that are targeted exclusively by clusters 0 and 2, respectively. Bacterial effector PBMs from pathogenic EPEC, *Shigella flexnerii, Legionella pneumophila* and *Citrobacter rodentium,* as well as resident *Aeromonas enteropelogenes, E. coli* (MS 115-1 and MS 198-1) and *Providencia stuartii* (ATCC 25827) strains, have a total of 38 interactions with these PDZ domains in the same affinity range. Comparison of binding energy differences (ΔΔG in kJ/mol) (Fig.2, d) reveals PBMs of Map from EPEC (`IADNVQDTRL`) and from *Citrobacter rodentium* (`IADCVQDTRL`) could compete with CFTR (`TEEEVQDTRL`) for interactions with the second PDZ domain of Na^+^/H^+^ exchange regulatory cofactor protein 2 (NHERF2), the only PDZ domain of sorting nexin cargo protein 27 (SNX27) and that of SH3 and multiple ankyrin repeats domain protein 1 (SHANK1), binding the first of these with higher affinity than CFTR, its strongest human PBM binder. NHERF2 is essential for regulation of ion transport and receptor signaling in epithelial cells^51^, interacts with EPEC effector Map both *in vitro* and *in cellula* via a PDZ-PBM interface and play a role in its virulence^27^. Interactions with NHERF1 and 2 anchor CFTR to the membrane (NHERF1) and inhibit CFTR mediated Cl^-^secretion at the apical plasma membrane of intestinal and airway epithelial cells (NHERF2)^50^. In the gut, hyperactivation of CFTR channel activity leads to secretory diarrhea^50^. In the lungs, CFTR activation depends on formation of a complex with NHERF1^51^ and its interaction with NHERF2 contributes to its apical retention^51^. Disruption of these interactions is associated with disease phenotypes observed in cystic fibrosis and provide ideal conditions for bacterial replication^50^. Altogether, our results suggest that effectors with similar PDZ targeting profiles as PBM-containing ionic transport proteins could be relevant to both gut and lung diseases. Additionally, given host cytoplasm Ca^2+^ levels are sensed by the T3SS tip and relayed to its base to increase effector secretion^52^, targeting Ca^2+^ transporters could play a role in effector translocation.

Overall, our results show effectors from resident microbiota and pathogenic bacterial strains have tailored C-terminal sequences, allowing them to interact with human PDZ domains with similar and sometimes higher binding affinities than the strongest human binders, potentially aiding in the hijack of cellular functions by taking advantage of the existing binding and functional specificities within the human PDZ domain family.

### From fragments to full-length proteins: bacterial PBMs interact with human PDZ-containing proteins in cell extracts

To assess if the quantified PDZ-PBM interactions occur between the PBMs and the corresponding full-length PDZ-containing human proteins, we selected 20 PBMs from effectors belonging to 13 different strains, covering the full range of affinities measured for PDZ-PBM pairs and representing all tested PBM classes and measured their interactions against full length proteins in cell extracts by native Holdup^55^ (Figures 3 and S13-14) (see Methods). Briefly, in native Holdup full-length proteins from cell extracts are incubated with bait PBMs until equilibrium and the unbound fraction is then quantified by LC-MS/MS.

**Figure 3.**
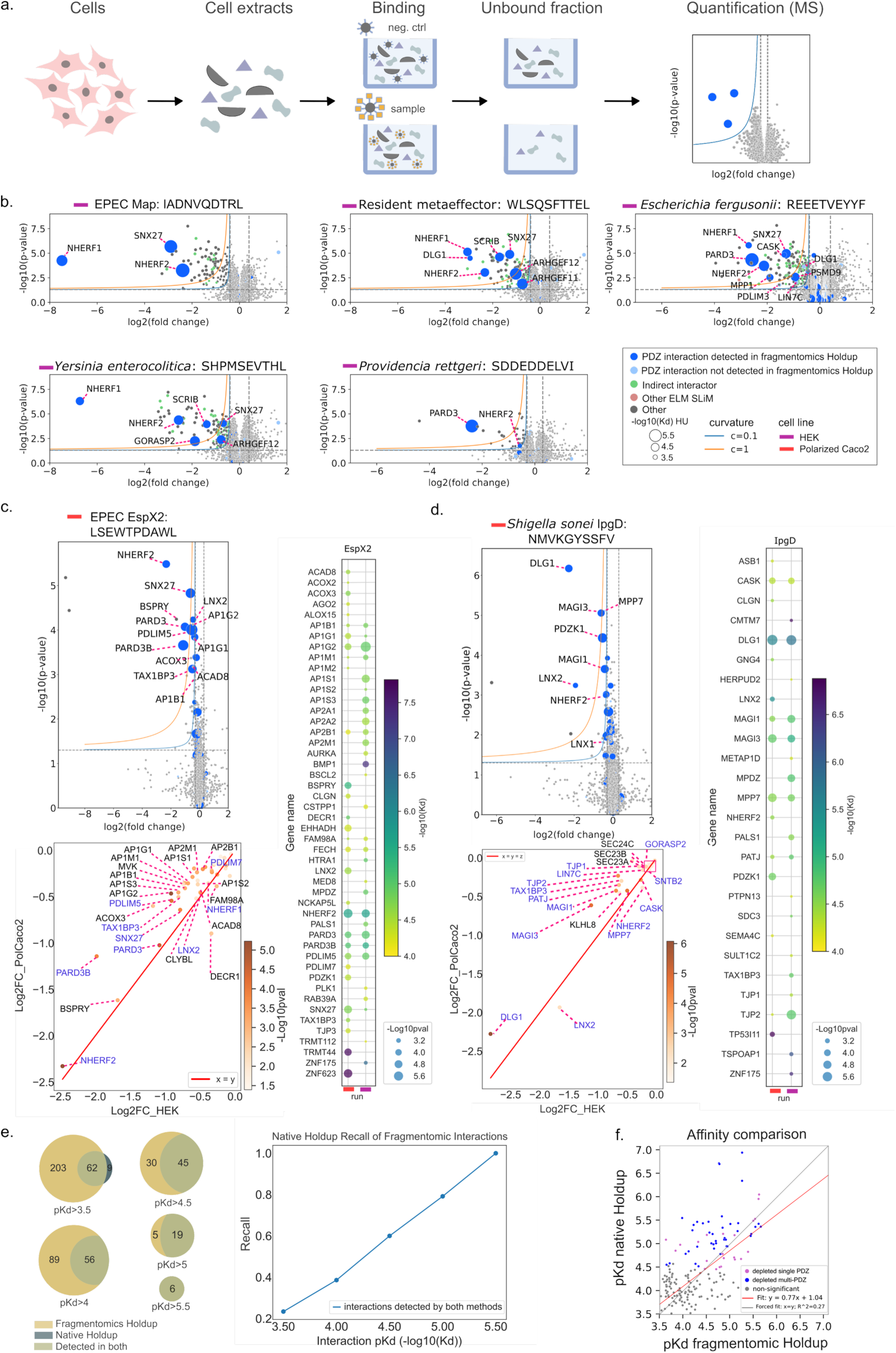
Native Holdup assay. **a,** Principle of the native Holdup for quantifying affinities with full-length proteins. **b,** Volcano plots with hyperbolic threshold curves showing log2 fold changes of MS-quantified proteins for 5/20 PBMs tested via native Holdup (MS with Orbitrap Exploris, ∼4500 proteins detected). Navy blue, sky blue, green, salmon and grey colored dots represent PDZ interactions detected by fragmentomic Holdup, PDZ interactions not detected by fragmentomic Holdup, indirect interactors of a PDZ binder detected by native Holdup (annotated from IntAct^43^ binary PPI data), proteins with Pfam^63^ domains that recognize a motif other than the PBM on the bait peptide, and depleted proteins not fitting into the previous groups, respectively. Significantly depleted PDZ protein gene names are annotated in each graph. **c,** (top left) Log2FC volcano plot for native Holdup for EPEC EspX2 PBM against polarized Caco2 cell extracts (see methods) with MS done by Orbitrap Astral (∼9-10k proteins detected), (bottom left) scatter plot of Log2FC of proteins significantly depleted (log2FC< -0.2, P-value <0.05) in both HEK293 and Caco2 cell extracts showing PDZ containing proteins in blue and others in black and (right) bubble plot showing apparent pKd for depleted interactors (log10pval>2.5, Log2FC<-0.2) EspX2 PBM against polarized Caco2 and HEK293 cell extracts. **d,** same as c) but for *Shigella sonei* IpgD PBM **e,** Venn diagrams (left) showing the overlap between significant interactions of fragmentomic and native Holdup experiments for different fragmentomic Holdup pKd thresholds, for all PDZ proteins quantified in the MS run (native Holdup data from the Orbitrap Exploris experiment) and plot (right) showing proportion of interactions quantified by both methods vs. fragmentomics Holdup pKd. **f,** Correlation between PBM affinities for PDZ domains (fragmentomic Holdup) and full-length PDZ-containing proteins (native Holdup; Orbitrap exploris experiment), showing non-significant (grey), depleted single-PDZ containing protein (pink) and depleted multi-PDZ containing protein (blue).

Of all interactions detected via fragmentomic Holdup assay for the 20 PBMs tested via native Holdup (See Methods), 265 correspond to PDZ domains in proteins that are expressed in HEK293 cells and that were reliably detected by MS. Of these, 62 are significantly depleted in native Holdup experiments with the corresponding full-length proteins, with recall by native Holdup increasing from ∼23 to 100% for increasing fragmentomic Holdup pKD thresholds (Figure 3e). The observed lower recall for weaker interactions is expected given the grey zone introduced earlier (Figure 1b). Only 9 interactions detected via native Holdup were not observed for the corresponding PDZ-PBM pairs by fragmentomic Holdup, indicating a low false positive rate and high specificity Comparison of affinities measured between PBMs and PDZ or the corresponding PDZ-containing full-length proteins shows a moderate correlation between the data (PCC=0.55), while a forced linear fit shows weak correlation (R^2^=0.27), with higher overall apparent affinities observed for interactions with full-length proteins (Figure 3f). This might partly be explained by the fact that multi-PDZ domain containing proteins can display avidity for a single ligand and thus increase the effective binding affinity (Figure 3f, Zambo et. al, 2022^55^).

For EPEC effector Map PBM, we recover previously reported interactions with NHERF1 and NHERF2, which are critical for its perturbation of cell surface projections27 and find an additional interaction with SNX27, an adaptor of the endosomal retromer complex implicated in endosome to plasma membrane protein recycling^56,57^, which has not been reported in the literature so far, indicating a potential targeting of the retromer complex by Map (Figure 3b).

Of seven class 2 PBM tested, 6 were shown to bind the partitioning defective 3 homolog (PARD3) protein, a master regulator of the Par polarity complex, including three from commensal effectors from *Escherichia fergusonii*, *Providencia rettgeri* (Figure 3b) and *Cedecea davisae* (Figure S14), two from *Legionella pneumophila* effectors (Figure S14c) and EPEC effector EspX2 (Figure 3c).

In subsequent native Holdup experiments we tested PBMs of effectors EspX2 and IpgD, from EPEC and *Shigella flexneri*, respectively, against HEK293 and polarized Caco2 cell extracts (see Methods). Since Caco2 are intestinal cells, they correspond to the bacterial tropism and might express relevant proteins not observed in HEK293 cells. The >2fold increase in proteome coverage (∼10,000 proteins) and the increased sensitivity (MS done with Orbitrap Astral) allowed probing the depleted region to greater depth, revealing PDZ containing interactors and additional binders (Figure 3c).

For EspX2 PBM, we quantified interactions with several PDZ-containing proteins, including some with known roles in membrane trafficking. Namely, SNX27 (retromer), as well as PDZ and LIM domain proteins 5 (PDLIM5) and 7 (PDLIM7), whose interactions with SNX17 via its C-terminal PBM play a role in Commander-dependent intracellular membrane trafficking at the endosomal compartment^58,59^. In the PDZ-PBM Holdup experiments (Figure 1), we detected interactions between EspX2 PBM with PDZ domains from PDLIM1,2,4,5 and 7, with mean Kds of 71µM, 12µM, 8µM, 6µM and 46µM, respectively. SNX17 C-terminal PBM binds them with mean Kds of 36.0µM, 23.4µM, 11.2µM, 25.3µM and 18.2µM^59^, respectively, indicating possible perturbation of SNX17 interactions with PDLIM proteins by EspX2 based on higher binding affinities for three of them (PDLIM2,4 and 5) for the bacterial effector. Additionally, several clathrin-associated adaptor protein complex members (AP1s and AP2s) (Figure 3c, left), which participate in intracellular transport pathways in the trans Golgi network/endosomes (AP-1) and plasma membrane (AP-2)^60^, were depleted in EspX2 PBM native Holdup experiments. Interestingly, EspX2 C-terminal harbors an additional motif (LIG_deltaCOPI_diTrp_1) known to bind with mid-micromolar affinities to the adaptor complexes medium subunit (PF00928), present in two of the observed binders: AP1M1 and AP2M1. It is likely that EspX2 binds them directly and that other AP proteins display binding specificity for this motif. While it is known that *Salmonella* effector SpvB perturbs membrane trafficking via clathrin and AP-1^61^, to our knowledge, this is the first time it is shown for EPEC, thus enriching our knowledge of the pathogen’s biology.

For IpgD, a well characterized effector with phosphatidylinositol 4,5 bisphosphate (PI4,5P2) 4-phosphatase activity implicated in membrane ruffling at entry site^62^, we quantify interactions between its PBM and several PDZ containing proteins, including several involved in tight-junction assembly and organization (Figure 3d). For instance, all three members of the stardust protein complex (MPP7-DLG1-LIN7C) were depleted for both HEK293 and polarized Caco2 cells. This complex plays a crucial role in apicobasal polarity and tight-junction maintenance^62^. Additionally, tight-junction proteins 1 and 2 (TJP1-2), PATJ and MPDZ, were also depleted and are all involved in cell junction organization. These interactions suggest that host cell membrane perturbation by IpgD might extend beyond known molecular mechanisms to include PDZ-containing proteins. We also detect interactions with non-PDZ containing protein Kelch-like protein 8 (KLHL8) a substrate adaptor for an E3 ubiquitin ligase complex involved in protein degradation, and transport proteins (SEC23A, SEC23B, SEC24C) discussed in the next section.

Altogether, the native Holdup results show that PBMs from resident and pathogenic bacteria can interact with full-length human PDZ containing proteins and sequence specificities observed for PBM clusters accurately reflect binding preferences for full-length partners. Further, bacterial PBMs can harbor motifs specific to different binding modules other than PDZ domains, as was observed for EspX2 interactions with AP proteins, hinting at a deeper layer of complexity in the molecular dialogue between bacteria and their host.

### The C-terminal PBM of NleA drives interactions involved in hijacking of transport pathways to remodel the host membrane

Next, we sought to validate interactions quantified via fragmentomic and native Holdup assays for one PBM, with the corresponding full-length effector. EPEC effector NleA and NleAΔPBM (deletion of last 10AA from C-terminal), as well as the PBM alone, were screened via native Holdup assay against extracts from HEK293, Caco2 and polarized Caco2 cells. Full-length NleA interactions with PDZ containing proteins have been explored in the past, allowing us to have a benchmark of positive hits^28^ (Figure S4).

NleA PBM interacts with 17 different PDZ containing proteins present in polarized Caco2 cell extracts (Figure 4a, left), 16 of which are also significantly depleted for HEK293 and Caco2 cell extracts (Figure 4b, left). Overall, we recover all previously identified NleA PDZ containing interactors in the literature and report new interactions with 11 PDZ containing proteins (Figure 4a, left and S4). Overall, between fragmentomic and native Holdup experiments, we recover (14/15) human-bacterial PDZ-PBM mediated PPIs reported in IntAct^43^.

**Figure 4.**
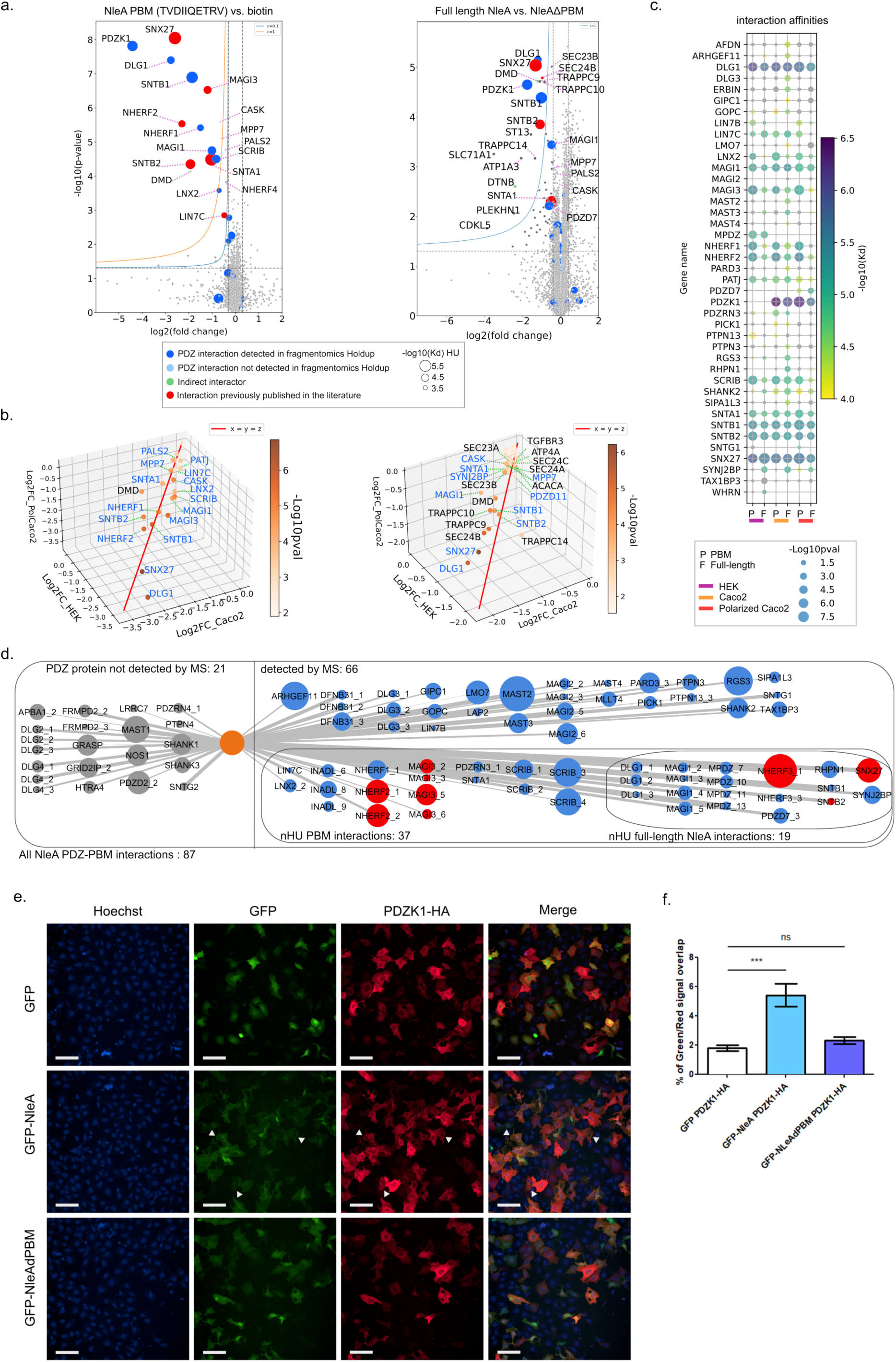
NleA: from fragments to full-length PPIs. **a,** Native Holdup Log2 fold-change MS volcano plots for proteins depleted from polarized Caco2 cell extracts using (left) NleA C-terminal PBM (`TVDIIQETRV`) vs. biotin and (right) full length NleA vs NleAΔPBM, respectively, as baits (for information about depleted proteins see legend). **b,** 3D scatter plots showing Log2(FC) for significant protein binders identified in HEK293, Caco2 and polarized Caco2 cell extracts for (left) NleA PBM (log2(FC) < -0.5, p value <0.05) and (right) full-length NleA (log2(FC) < -0.2, p value <0.05). In both scatter plots PDZ containing proteins are annotated in blue and non-PDZ containing protein in black **c,** pKd bubble-plot for all PDZ containing proteins detected in all runs for extracts from different cell lines/conditions for both NleA PBM (x-axis tick P) and full length NleA (x-axis tick F). **d,** Full PDZ-PBM PPI network for NleA PBM (TVDIIQETRV), showing PDZ-PBM fragmentomic Holdup interactions for proteins not detected by MS (grey) and insets showing proteins for which interactions were detected by HU but not native Holdup, detected by Holdup and native Holdup for the PBM and detected by native Holdup for full length NleA. **e,** U2OS cells co-transfected with pcDNA5-PDZK1-HA and pLVX-GFP, pLVX-GFP-NleA or pLVX-GFP-NleAΔPBM were stained with an anti-HA antibody (red) and Hoechst 33258 (blue). Triangles highlight cells were the green (GFP) and red (PDZK1-HA) signals overlap. Scale bar = 100 µm. **f,** Percentage of Green/Red signal overlap from images acquired in e.). n.s.: non-significant, ***P<0.001, one-way ANOVA, Dunnett’s multiple comparison test. N=4

Interestingly, certain interactions were unique to specific cell lines because the corresponding human proteins were not detected by MS for all cell lines, such as PDZK1 (only detected for Caco2 and polarized Caco2 cells), MPDZ (HEK293), PDZD7 (polarized Caco2) and LIN7B (HEK293, polarized Caco2) (Figure 4c). Among these, PDZD7 shows high RNA expression in GI tract tissues^47^, hinting at a possible role for PDZD7 during EPEC infection.

Of 19 interactions detected for NleA PBM via native Holdup with PDZ containing human proteins, 10 of them were also depleted for full length NleA vs. NleAΔPBM. Because the effective concentration of full length NleA on the beads is lower (∼2x) than that of the PBM, its capacity to deplete binding partners in the weaker affinity range diminishes, decreasing the percent recovery. However, equilibrium binding equations allow us to calculate the corresponding pKds for different bait concentrations and comparing them with those for the PBM (Figure 4c), showing good agreement between both measurement modalities.

When comparing the depletion profiles (Log2(FC) < -0.2 and P-value <0.05) for full length NleA vs. NleAΔPBM runs across the three different conditions (HEK293, Caco2 and polarized Caco2), we observe interactions with 10 PDZ containing proteins as well as several other interactors (Figure 4b). This confirms that the C-terminal PBM drives the interaction between the full-length human proteins and bacterial effectors, indicating PDZ-PBM interactions recapitulate the association for a significant portion of the corresponding full-length proteins and the native Holdup assay reproducibly quantifies them across different cell lines.

Among non-PDZ containing proteins, we detected interactions between full-length NleA and SEC24 A-C, which have been previously shown to be essential for localization of NleA and its associated virulence by inhibiting COPII-mediated vesicle transport^64–66^, as well as new interactions with trafficking protein particle complex II subunits 9 (TRAPPC9),10 (TRAPPC10) and 14 (TRAPPC14), potassium-transporting ATPase alpha chain 1 (ATP4A) and transforming growth factor beta receptor type 3 (TGFBR3).

Functional enrichment analysis of proteins significantly depleted by full-length NleA in all cell extracts (Figure 4b, right) against their common background (Figure S15a), shows significant enrichment of COPII-coated vesicle cargo loading and vesicle budding from membrane related processes (Figure S15b). Interestingly, SEC24 A-C along with SEC23 A-B drive the first annotation term exclusively and the second annotation term jointly TRAPPC9 and 10. The TRAPPII complex has been shown to bind to the COPI vesicle coat but not to COPII vesicles^67^ and we found no known interactions between TRAPPC9,10 and 14 with depleted SEC24 or SEC23 proteins, suggesting that NleA directly interacts with TRAPPII complex members in a C-terminal dependent manner. This implies that on top of the known perturbation of anterograde ER to Golgi transport via COPII inhibition, NleA might also disrupt intra-Golgi and early endosome to late Golgi retrograde traffic via perturbation of TRAPPII complex-associated membrane tethering of coated-vesicles^67^. Recent work demonstrated that NleA undergoes host-mediated mucin-type-O-linked glycosylation^68^ and that interaction with SEC24 plays a role in it, but the exact molecular mechanisms remain unknown^69^. The observed interactions for NleA with TRAPPII complex proteins suggest they might also play a role in its glycosylation, given their involvement in intra-Golgi processes.

Among depleted PDZ-containing interactors, earlier work showed SNX27 mediates recycling of beta-2 adrenergic receptor (ADRB2) from the early endosome to the plasma membrane, via an interaction between its PDZ domains and ADRB2’s C- terminal PBM^57^ and regulates apical targeting of mucin MUC17^70^, suggesting that NleA’s interaction with SNX27 via its PDZ domain could lead to impaired protein recycling to the plasma membrane, as has also been observed for other bacterial pathogens^71^.

Besides protein transport, we observed enrichment for several ionic transport processes (Figure S15) driven by PDZ containing proteins DLG1, SNTA1, CASK PDZD11 and PDZK1 (exclusively for Caco2 cells), as well as the potassium-transporting ATPase alpha chain 1 (ATP4A), which does not contain a PDZ domain. Among these, PDZD11 has been shown to bind to all plasma membrane Ca^2+^ATPase b-splice forms and to regulate their localization^72,73^. ATP4A, which was mildly depleted in all native Holdup experiments with full length NleA, is a catalytic subunit of the H^+^/K^+^ ATPase pump that exports H^+^ ions and imports K^+^ ions across apical membranes, creating a proton gradient in the gastric lumen which acidifies gastric juices down to pH 1. Perturbing electrolyte balance across apical membranes could have the dual role of affecting cell polarity as well as creating favorable extracellular conditions for bacteria.

TGFBR3 regulates the actin cytoskeleton and promotes activation of CDC42 and inhibition of NF-κB through interactions with beta-arrestin 2 (ARRB2)^74^; interestingly, CDC42 is a known target of EPEC Map effector through its interaction with F-actin^75^ and plays a crucial role in EPEC-induced membrane remodeling. Further, EPEC effector NleH inhibits NF-κB in a PDZ-binding dependent manner via its C-terminal PBM^29^. Thus, NleA may be another effector that targets CDC42 and regulates NF-κB indirectly through TGFBR3.

Finally, dystrophin-associated glycoprotein complex members SNTA1, SNTB1, SNTB2 and DMD, were consistently and significantly depleted across native Holdup experiments for both full-length NleA and its PBM, along with DTNB, which anchors SNTA1 and DMD to basal membranes. Targeting of syntrophin proteins has been observed for EPEC NleA^28^ and *Salmonella enterica* effector SseM^33^, via their respective C-terminal PBMs, suggesting a convergent targeting of the complex by bacteria with different cellular lifestyles. Previous work demonstrated that SseM interacts with α-dystrobrevin (DTNA) indirectly via SNTB2^33^, suggesting that depleted non-PDZ containing members of the glycoprotein complex might be “dragged” by direct PDZ containing interactors.

The strongest identified interaction both for the PBM and for full length NleA, are with PDZK1 (the first PDZ domain and the full-length protein, respectively), a scaffold protein that stably localizes transmembrane mucin MUC17 to enterocyte apical membranes in the small intestine, via interaction of MUC17 C-terminal PBM with the first PDZ domain of PDZK1^76^. In mice, absence of Pdzk1 results in intracellular accumulation of Muc3 (MUC17 mouse orthologue)^77^, suggesting that direct binding competition between NleA and MUC17 for PDZK1 could lead to a depletion of MUC17 at apical membranes, potentially thinning the glycocalyx to facilitate bacterial adhesion to the outer membrane. Co-transfection of PDZK1 and NleA into U2OS cells, shows a significant difference in colocalization between NleA and the negative control, while the NleAΔPBM shows no difference (Figure 4e-f), confirming that the interaction between PDZK1 and NleA takes place inside the cell in a PBM-dependent manner.

Overall, our interaction screens for NleA provide a richer picture of the molecular mechanisms perturbed by EPEC during infection. We provide evidence of both PDZ and non-PDZ containing proteins involved in its known perturbation of protein trafficking, ionic transport processes and cell-cell anchoring and adhesion. Altogether, these perturbations could lead to a cell surface remodeling, electrolyte imbalance across apical membranes and perturbation of cellular adhesion, which might benefit bacterial access to the cell surface during infection. Remarkably, all these interactions are mediated by a 10 amino acid C-terminal SLiM, that targets human proteins with different characteristics and functions.

### Additional C-terminal overlapping SLiMs expand complexity of the host-microbe molecular dialogue

Protein multifunctionality is a known feature of the human proteome^78^ and can be mediated by post-translational modifications that reshape molecular interactions^79–81^. Previous work has shown phosphorylation substantially alters PBM interactions with PDZ domains, modulating their affinity and specificity^82^. Viral oncoprotein HPV16-E6, binds either PDZ or 14-3-3 domains in a phosphorylation dependent manner^81,83^ and *Pseudomonas syringae* T3SS effector HopQ1 gets phosphorylated in plant cells and interacts with 14-3-3 proteins, possibly affecting plant immunity^84^. Effector phosphorylation is also known to occur inside human cells and to play an important role in bacterial virulence^5^.

In addition to harboring PBMs, overlapping motifs occur on the C-terminal of most effectors (172/201) for which we measured PDZ-PBM interactions, including targeting, docking, ligand, modification, degradation and cleavage motifs (Figure 5a).

**Figure 5.**
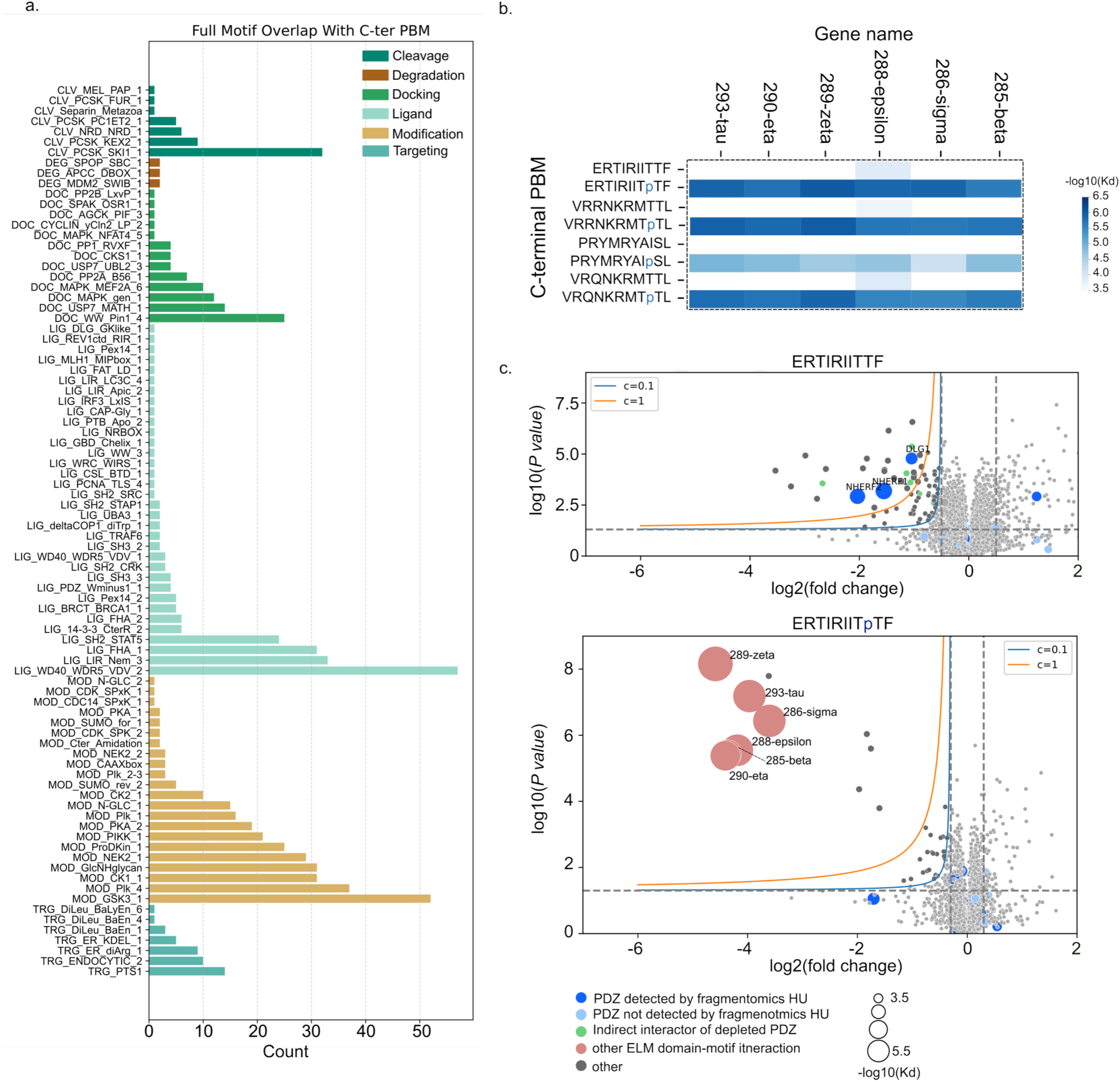
Analysis of overlapping C-terminal motifs. **a,** Counts of motifs reported in ELM^40^ that overlap with C-terminal PBMs of tested bacterial effectors. **b,** Domain-motif binding affinities (-log10(Kd)) measured by Holdup assay for 14-3-3 domains with C-terminal PBMs and their phosphorylated versions (pPBMs), for motifs matching the LIG_14-3-3_CterR_2 regular expression profile, and **c,** native Holdup depletion profiles for a selected PBM (`ERTIRITTF`, top) and its phosphorylated version (`ERTIRITpTF`, bottom) with PDZ domain containing proteins colored blue and 14-3-3 proteins colored salmon.

To test whether these could be functionally relevant, we synthesized phosphorylated versions of PBMs matching the LIG_14-3-3_CterR_2 motif and quantified the interactions of 6/7 14-3-3 proteins against the regular PBM and the phosphorylated (pPBM) version by fragmentomic Holdup (Figure 5b). All 14-3-3 proteins were found to bind with similar affinities all pPBMs while interactions with their PBM counterparts could not be detected (Figure 5b). To assess if these interactions occur with the corresponding full-length proteins, we tested one PBM-pPBM pair, namely `ERTIRITTF-ERTIRITpTF`, by native Holdup (Figure 5c). For the regular PBM, interactions with NHERFs 1 and 2 and DLG1 were detected (Figure 5c top), while for the pPBM all interactions detected with 14-3-3 proteins by Holdup were also significantly depleted by native Holdup (Figure 5c, bottom), while the interactions with PDZ domain-containing proteins were not detectable.

Bacterial effector functions mediated by disordered regions could therefore be more complex than currently understood, with short sequence stretches potentially encoding multiple functions that could depend on host cell state and context.

## Discussion

Protein-protein interactomes provide a systems view of biological function. By quantifying their associated binding affinities, we develop a richer understanding of molecular mechanisms. Thus, we charted the largest quantitative interactome to date between human PDZ domains and bacterial C-terminal PDZ-binding motifs, increasing the number of reported affinities between human domains and bacterial SLiMs by a factor of ∼200, compared to current literature.

We show pervasive targeting of human PDZ domains by effectors from both the gut microbiota as well as gut and lung pathogens. Further, PBMs from pathogens and resident gut strains cluster together with human PBMs based on affinity profiles for common PDZ targets and in several instances, bind them with similar or higher affinity than the strongest human binder. Such cases were observed for effectors from well-known pathogens (Map, NleA, EspH, IpgD) as well as effectors from resident gut strains with uncharacterized functions. Interestingly, members of these clusters have well defined C-terminal amino acid compositions and are enriched for common and unique human cellular functions, hinting at a function-directed evolutionary tailoring of human protein sequences that drives binding specificities and affinities within the PDZ domain family, a phenomenon exploitable by bacterial effectors. Of note, functions linked to cell polarity, tight junction assembly, protein localization to cell periphery and ion transport, were enriched for most clusters, are known to be perturbed by pathogens and could play a role in disease associated with gut dysbiosis.

Our native Holdup experiments show that interactions between fragments (PDZ-PBM) recapitulate a significant portion of the corresponding interactions with full-length human proteins and accurately reflect sequence-based PDZ binding preferences. Members (DLG1, SCRIB, PARD3, PATJ) of all three human cell polarity complexes (SCRIB-DLG1-LGL^45^, PAR3/PAR6/aPKC and CRUMBS/PALS1/PATJ) were depleted for several PBMs, indicating possible perturbation of cell polarity by pathogens and resident strains alike. Similarly NHERF 1, 2 and 3 are targeted broadly, in both gut and lung tissues, where their interactions with PBM-containing proteins are crucial for their roles in fluid and electrolyte balance, both of which are perturbed by gut and lung pathogens and relevant for virulence. Additional native Holdup experiments with different cell-lines, revealed targeting of transport related PDZ-containing and non-PDZ-containing proteins for EPEC effectors EspX2 and NleA. Namely, full-length NleA is shown to interact in a C-terminal dependent manner with proteins involved in COPII-mediated ER to Golgi anterograde transport (SEC24 A-C, SEC23 A-B), COPI-meditated intra-Golgi and early endosome to late Golgi retrograde traffic (TRAPPC9,10 and 14). EspX2 PBM interacts with PDLIM5 and 7, which are involved in membrane trafficking at the endosomal compartment as well as several members of adaptor protein complexes 1 and 2, involved in transport pathways in the trans Golgi network/endosomes (AP-1) and plasma membrane (AP-2)^60^. Additionally, these and several other effectors interact with SNX27, potentially influencing recycling of membrane proteins by the retromer complex. Overall, our results extend previous knowledge of perturbation of COPII mediated transport by pathogenic *E. coli*, to include COPI, AP-complexes and the retromer, depicting a fuller picture of the perturbation of intra-cellular protein transport by the pathogen. Combined with the implication of targeted PDZ containing proteins in the localization of proteins to the cell periphery and their stabilization at membranes (e.g. PDZK1 for MUC17), our results suggest a global hijacking of protein transport from the ER all the way to the membrane, as well as a possible alteration of membrane protein composition. We argue this constitutes plausible evidence for glycocalyx remodeling by EPEC and warrants further investigation.

Finally, the existence of overlapping C-terminal motifs targeting other human protein modules indicates an additional layer of complexity exists in the molecular dialogue between human cells and secreted bacterial effectors.

Owing to the modular nature of the human proteome, systematically quantifiying interaction affinities between human protein interaction modules and bacterial effector modules that target them, will provide insight into competition mechanisms at the molecular level, potentially paving the way towards novel therapeutic strategies in the case of pathogens and a better understanding of molecular mechanisms implicated in disease for resident microbiota bacteria.

Altogether, we show the potential that a quantitative interactome between protein fragments has for the elucidation of effector protein function. While we only explored a small subset of our interactome for validation between full-length effectors and human proteins, this exploration yielded rich insight into perturbation of known biological functions and provided evidence to posit new hypotheses related to the identified interactions. As such, the data presented here is a rich resource for the community and provides new avenues for discovery in the field of host-microbe interactions.

## Materials and Methods

### Bacterial culture for PDZ production

The PDZOme V.2 library, consisting of all 266 human PDZ domains as His-MBP-PDZ constructs, was prepared as previously described^85^.

Briefly, PDZ domains were expressed in *E. coli* BL21 (DE3) overnight at 25°C in autoinduction media, in a culture volume of 100ml in 500ml flasks or in 24ml (12x2 ml/well) in 24 well deep well plates^86^. Cultures were grown to an OD of 8-12, centrifuged and resuspended in 50ml (for 100ml cultures) and 6ml (for 24ml cultures), of lysis buffer (300mM NaCl, 50mM Tris-HCl, pH 7.5, 1% Triton X-100, 10% glycerol and trace amounts of Lysozyme) and frozen at -20C.

Two copies of the PDZome were purified by sequential double affinity chromatography (DAPF), and by single affinity chromatography (SAPF), respectively.

### Double affinity purification (DAPF)

Frozen pellets were thawed at 37°C for 10min, resuspended after adding trace amounts of DNAse and MgCl2 and sonicated at 40% power for 3min using 30s on/off cycles. The resuspended pellets were then centrifuged at 12,000g for 10min and the supernatant was recovered in 50ml falcon tubes.

Purification of supernatant lysates follows the double affinity purification protocol from^41^. Briefly, supernatant lysates were loaded into prepacked 5ml high flow His-Trap FF crude Ni columns (Cytiva) in buffer A (300mM NaCl, 50mM Tris-HCl, 10mM Imidazole, pH 7.5) at a rate of 5ml/min, washed in 10 column volumes of wash buffer (300mM NaCl, 50mM Tris-HCl, 50mM Imidazole, pH 7.5) and eluted in ∼17ml of elution buffer 1 (300mM NaCl, 50mM Tris-HCl, pH 7.5, 250mM Imidazole). The Ni elution was directly loaded onto prepacked amylose resin columns (Cytiva, high flow), washed with 10 column volumes of buffer A’ (50ml of 150mM NaCl, 25mM Hepes, 10% glycerol at pH 7.5) and eluted in ∼4ml of elution buffer 2 (150mM NaCl, 25mM Hepes, 5mM maltose, 10% glycerol, 1mM TCEP, pH 7.5). Purified PDZ domains were frozen at -20°C. Prior to freezing, PDZ quality was assessed by SDS-page gels and capillary electrophoresis (LabChip GX) and concentration was calculated from UV absorbance measured on PHERAstar FSX (BMG Labtech, Offenburg, Germany) microplate reader with the 295 ± 10 nm–350 ± 10 nm (Tryptophan-fluorescence) band-pass filter. A fraction of the concentrated solutions was diluted to get 2x5ml aliquots at 4µM to be frozen separately.

All DAPF purifications were performed on AKTAxpress (Cytiva) systems, with a small fraction done manually on the benchtop.

### Single Affinity Purification (SAPF)^85^

Thawing, re-suspension and sonication were performed as in DAPF, except in 24 well DW plates with an additional pooling step (12x500µl pooled into a single well for 6ml lysate/target). Lysates were purified on Freedom EVO 200 automated liquid handling system (TECAN). First, Ni beads (Cytiva, Sepharose fast flow chelating beads) were equilibrated in buffer A (300mM NaCl, 50mM Tris-HCl, 10mM Imidazole, pH 7.5) in a solution containing 50/50 v/v dry beads/buffer A, then loaded into each well of the 24 well plate with the 96 tip automated pipetting head, for a total equivalent volume of 200µl dry Ni beads/well. Binding to the Ni beads was performed under mixing with 800µl aspirate/dispense cycles for 15min. The Ni bead-bound lysates were then transferred into a 96 well filter plate (35 micron pores, Macherey-Nagel) and placed on a vacuum block, where the liquid unbound fraction was vacuumed. The bound fraction was washed twice in buffer A (1ml/well x 2), then washed a single time in wash buffer (300mM NaCl, 50mM Tris-HCl, 50mM Imidazole, pH 7.5) before elution in 2.5 ml buffer B (300mM NaCl, 50mM Tris-HCl, pH 7.5, 250mM Imidazole, 5mM dithiothreitol). Concentration of the purified PDZ was calculated from UV absorbance as for DAPF, then each one was diluted to 4µM and stored as 4x1-2ml aliquots at -20°C. Prior to freezing, PDZ quality was assessed by SDS-page gels and capillary electrophoresis (LabChip GX).

### Reagents for crude peptide array

All solvents and chemicals were used without any further purification and purchased as peptide grade when available. N-methylpyrrolidinone was purchased from Biosolve. isopropylcarbodiimide and collidine were purchased from Sigma. The following L-amino acids were used: Fmoc-Ala, Fmoc-Arg(pbf)-OH, Fmoc-Asn(Trt)-OH, Fmoc-Asp(Otbu)-OH, Fmoc-Gln(Trt)-OH, Fmoc-Glu(Otbu)-OH, Fmoc-Glu-Otbu, Fmoc-Gly-OH, Fmoc-His(Trt)-OH, Fmoc-Ile-OH, Boc-Ile-OH, Fmoc-Leu-OH, Fmoc-Val-OH, Fmoc-Lys(boc)-OH, Fmoc-Ser(Otbu)-OH, Fmoc-Thr(Otbu)-OH, Fmoc-Tyr(Otbu)-OH, FmocTrp(Boc)-OH, Fmoc-Phe-OH, and they were purchased from either Protein Technologies, Novabiochem Merck or IRIS Biotech (Germany). Biotin was purchased from Sigma. Fmocpreloaded resin was used and purchased from Merck Millipore.Fmoc-8-amino-3,6-dioxaoctanoic acid (Fmoc-Ado-OH) was purchased from Flamma Group, Italy.

### Synthesis of crude peptide array

Crude peptide libraries were synthesized using a parallel 96-well format with the Intavis Multipep RSi. We used filter plates from NUNC (Thermo Fisher) and loaded each well with 10 mg of Fmoc-amino acid preloaded polystyrene resin (Merck Millipore). Fmoc-amino acids (Fmoc-AA) were diluted to 0.3 M in dimethylformamide (DMF) with 0.3 M OxymaPure.

We performed three consecutive peptide couplings using 100 μl of Fmoc-AA and 11 μl of DIC solution (3 M in DMF), with coupling times of 5, 15, and 60 minutes. To remove the Fmoc group, we washed the wells twice with 120 μl of 25% piperidine in DMF for 2 and 8 minutes. Washing was done by adding 150 μl DMF to each well five times. For biotin coupling, we used a 0.3 M solution in DMSO with 0.3 M OxymaPure, activated by DIC, with extended coupling times of 15, 45, and 360 minutes. To cleave the peptides, we added a total volume of 1 ml of a TFA solution (93% TFA, 3% triisopropylsilane, 1% thioanisole, 3% H2O) to each well for 2 hours. The TFA was added in portions of 5x100 μl separated by 1 min. Then after 30 min 2x150 μl and then after additional 1.5 hour 3x100 μl. After TFA addition, we reduced the volume to about 150 μl in the deep well collector plate and precipitated the peptides by adding diethyl ether.

The peptides were transferred to Waters Solvinert plates and thoroughly washed five times with diethyl ether using a 8-pin manifold. The volume was approximately 500 µL to each well. A small fraction of the peptide slurry was then added to a microtiter plate and dried, followed by UPLC-MS analysis after solubilization in DMF/H2O (1:1). We used a Waters Acquity UPLC system equipped with a TUV detector at 214 nm and a LCT Premier XE mass spectrometer from Micromass for purity and MS identity analysis. The buffers used for the analysis were 0.05% TFA in water (Buffer A) and 0.05% TFA in acetonitrile (Buffer B), employing a gradient from 5% to 60% Buffer B over 3.5 minutes on a Waters BEH C18 column (1.7 μm, 2.1 mm x 50 mm) at a flow rate of 0.45 ml/min. Internal standards for fluorescent Holdup normalization were also synthesized as crude peptides. No significant differences were observed between the HPLC-purified and crude peptides during the Holdup experiments.

We confirmed the predicted peptide masses through mass spectrometry, with most peptides showing a purity level exceeding 80%. Average peptide concentrations were calculated based on the total dry weight of the entire 96-well plate, and we used a molar excess of 10-50 times (based on dry weight) in Holdup experiments to accommodate variability in peptide yield.

### Peptide solubilization

A simple heuristic was used for the solubilization of peptides based on positively charged, negatively charged and hydrophobic residue content. In general, positively charged peptides were solubilized in 1% NaHCO3 solution and negatively charged peptides in 5% Acetic acid. Hydrophobic peptides were initially tested for solubility in DI water, with only a handful not solubilizing. The confirmed hydrophobic peptides were lyophilized overnight to remove water and then solubilized in either DMSO or DMF, the latter for thiol-containing residues. Solubilization volumes were calculated based on the dry mass of the peptide powders and their molecular weights, for a final concentration of 10mM. Four sets of mother plates of the 4x384 peptides at 10mM were distributed onto 96 well V-shape plates and stored at -20°C.

### Holdup assay resin pre-saturation

Streptavidin coated sepharose beads (Streptavidin Sepharose High Performance, Cytiva) were washed twice in DI water and twice in HU buffer (50 mM TRIS pH 7.5, 300 mM NaCl, 1 mM TCEP) by centrifugation in 15ml Eppendorf tubes for 5min at 500g.

A solution of 40ml beads (1/4 "dry" beads v/v) in HU buffer was put in a trough, thoroughly mixed by pipetting up and down with the 96-tip automated head (Tecan Liquid Handling System) and 80µl were dispensed into each well of 4x96 V-shape plates with the Tecan 96 tip head.

PBM plates for the assay were prepared by diluting the mother (10mM) plates as follows: 180µl final volume, 1.98µl PBM at 10mM (diluted ∼22 times), for a final concentration of 110µM. These daughter plates were used for resin saturation, as described below.

80µl of PBM at 110µM were dispensed with an automated 96 tip head (Freedom Evoware, Tecan) into each well of the 4x96 V-shape well plates (Greiner) containing the ¼ v/v streptavidin bead solution, for a final concentration of PBM of 55µM and a final bead content of 12.5% by volume. This mix was incubated on a vibrating plate at 1200 rpm (Tecan T-shake) for 2 hours.

### Holdup plate preparation

40µl of the above pre-saturated resin solution was dispensed into each well of a 384 well filter plate (Merck, MZHV, 384 filter plate) for a final dry bead volume of 5µl/well.

The wells were washed with 10 resin volume HU buffer before depletion with 25µl of 1mM biotin for 15min under agitation (t-shake, 1200rpm). The plates were then washed with 100µl HU buffer, vacuumed and either used immediately (see next section) or topped with 20% ethanol, plastic filmed and stored at 4°C for later use.

### Holdup Interaction assay

Purified His6-MBP-PDZ domains from either SAPF or DAPF purifications were thawed in water at room temperature for 5min. The SAPF PDZ had been previously frozen in several 2ml aliquots at 4μM, whereas the DAPF PDZ had been either aliquoted as 2x5ml at 4μM or stored concentrated in 2ml aliquots following elution from the amylose resin. In the latter DAPF case, concentration was determined before freezing and corroborated after thawing by UV absorbance at 280nm, then diluted accordingly for a minimum volume of 5ml at 4μM. All PDZ solutions were supplemented with 50nM fluorescein and 100nM m-Cherry for signal normalization.

Once the 384 filter plates (MZHV) containing the PBM saturated beads were washed with HU buffer after biotin depletion and vacuumed, they were centrifuged at 700g for 2min before loading 10 µl of PDZ @4µM into each well of the 384 plate (either manually with the automated multi P-30 or on the Tecan with the needles of the liquid handling system). A 384 round-bottom well black receiving plate (Corning) was placed under the filter plate during PDZ loading and incubation. The plate was incubated until equilibrium (15 min), then covered with its transparent plastic lid, taped along the seams to the 384 well Corning read-out plate and centrifuged at 700g for 2min. Finally, the recovery plate was read on a PHERAstar FSX (BMG Labtech, Offenburg, Germany) microplate reader by using 485 ± 10 nm–528 ± 10 nm (fluorescein), 575 ± 10 nm–620 ± 10 nm (mCherry), and 295 ± 10 nm–350 ± 10 nm (Trp-fluorescence) band-pass filters, respectively.

Holdup plates were recycled using the protocol described in Gogl et. al^41^ and used up to ∼20 times.

The assay itself was run with replicates throughout the experimental campaign, with multiple measurements performed for 44 PDZ domains for a total 8831 binary PDZ-PBM interactions that were measured 23037 times.

### Native Holdup Interaction assay

#### Cell culture for native Holdup

For native Holdup experiments at the IGBMC, HEK293 cell cultures and extracts were prepared as in^55^.

For native Holdup runs at the CRCM, HEK293 cells (ATCC CRL-1573) were cultured in six T175 flasks (Falcon) containing DMEM + GlutaMAX + 4.5 g/L glucose + pyruvate medium (Gibco) supplemented with 10% fetal bovine serum (FBS), at 37°C in a humidified atmosphere containing 5% CO₂. Cells were grown until confluence and used to prepare extracts (see next section).

Human colon Caco-2 cells (ATCC HTB-37) were maintained in DMEM supplemented with 10% FBS, 1% sodium pyruvate, 1% non-essential amino acids, and 1% penicillin-streptomycin. Cells were seeded in T-175 flasks. To induce differentiation, cells were cultured for 21 days, with the culture medium being changed three times per week. Total cellular extracts were prepared at two time points: after 5 days of culture, when cells had reached confluence, and after 21 days of culture when the cells were differentiated.

### Cell extracts

HEK293, Caco2 and polarized Caco2 cells were grown to confluency, washed in 1x with PBS and collected by scraping with 5-10ml ice-cold lysis buffer (50 mM Hepes-KOH, pH 7.5, 150 mM NaCl, 1% Triton X-100, 5× complete EDTA-free protease inhibitor cocktail 5x, 5 mM TCEP, and 10% glycerol), sonicated in 2ml tubes (Starsted, ref. 72.694) for 4x20s with 1s pulses at 20% amplitude on ice, then put on the spinning wheel at 4°C for 30min, before being centrifuged at 12,000 rpm at 4°C for 20min. Extract concentration was determined by BCA and extracts were flash frozen at 1.5, 2 or 6mg/ml as 120µl aliquots in 0.5ml Protein LoBind tubes (Eppendorf) and stored at -80°C.

### Resin preparation and native Holdup experiment

Resin saturation and incubation were performed as in^55^. Briefly, streptavidin saturated sepharose beads (Streptavidin Sepharose High Performance, Cytiva) were washed twice in DI water and thrice in HU buffer (50 mM TRIS pH 7.5, 300 mM NaCl, 1 mM TCEP) for equilibration. For resin saturation with peptides or biotin, the equilibrated resin solution was diluted to 1/6.7 v/v content of dry resin, and 167 µL of this mix were incubated with 8.3 µL of peptide or biotin at 50 µM for 1h at 5°C on a tuberoller or a spinning wheel. After saturation, the resins were washed once with 400 µL of HU buffer and depleted with a 100 µM biotin solution in 1 mL of HU buffer for 20min at room temperature. After biotin depletion, the resins were washed three times in 400 µL of HU buffer.

For incubation with cell extracts, the pre-saturated resin solutions were centrifuged for 15s at 2,000g, the supernatant was discarded and 100 µL of cell lysate at 1mg/ml was added to each tube and incubated with the bait-bound resin for 1.5h at 4°C.

After incubation, the resin was centrifuged for 15s at 2,000g and 50 µL of supernatant (top half) were immediately collected into a second tube, centrifuged again for 15s at 2,000g and half of the supernatant (25 µL) was collected and frozen at -20°C immediately after collection for analysis by mass spectrometry (MS).

### Reagents for MS (IGBMC)

The following reagents for MS were purchased from Sigma Aldrich: Acetonitrile MS grade (ACN), Formic acid (FA), Trifluoroacetic acid (TFA), Trichloroacetic acid (TCA), Tris-HCl, Iodoacetamide, Urea.

### Liquid digestion (IGBMC)

Protein mixtures were TCA-precipitated overnight at 4°C, pellets were washed twice with 1 mL cold acetone, dried and dissolved in 2 M urea in 0.1 mM Tris-HCl pH 8.5 for reduction (5 mM TCEP, 30 min.), and alkylation (10 mM iodoacetamide, 30 min.). Trypsin digestion was carried out at 37°C and overnight. Peptide mixtures were then desalted on C18 spin-column and dried on Speed-Vacuum before LC-MS/MS analysis.

### MS analysis (IGBMC)

Samples were resuspended in 0.1%TFA and then analyzed using an Ultimate 3000 nano-RSLC (Thermo Scientific, San Jose California) coupled in line with a quadrupole - orbitrap Exploris 480 via a nano-electrospray ionization source (Thermo Scientific, San Jose California) and the FAIMS pro interface. Less than 200 ng tryptic peptides were loaded on a C18 Acclaim PepMap100 pre-column (300 µm ID x 5 mm, 5 µm, 100Å, Thermo Fisher Scientific) for 1 minutes at 15 µL/min with 2% ACN, 0.1% FA in H2O and then separated on a C18 PepMap nano-column (75 µm ID x 25 cm, 2.6 µm, 150Å, Thermo Fisher Scientific) with a 70 minutes linear gradient from 7% to 35% buffer B (A: 0.1% FA in H2O; B: 0.1% FA in 80% CAN) followed by a regeneration step at 90% B and a equilibration at 7% B. The total chromatography was 95 minutes at 450 nl/min and the oven set to 45°C. The mass spectrometer was operated in positive ionization mode in Data-Dependent Acquisition (DDA) for two FAIMS compensation voltages (CV = -45V & -55V). The DDA cycle consisted of one survey scan or MS1 (330-1400 m/z, 90,000 FWHM) followed by MS2 spectra (HCD; 30% normalized energy; 8 m/z window; Scan Range 140-1400 m/z; 22,500 FWMH). The cycle time was limited at 1.5 sec for the CV -45 and 1.2 sec for CV -55. The Automatic Gain Control (AGC target) were set to 3E6 and 1E5 respectively for MS1 and MS2 scans, and the maximum injection time (IT) was set to 50 ms for both scan modes. Unassigned and single charged states were rejected. The exclusion duration was set for 30s with mass width was ± 10 ppm.

### MS data processing (IGBMC)

Proteins were identified with Proteome Discoverer 2.5 software (Thermo Fisher Scientific) and *Homo Sapiens* proteome database (SwissProt, release 2024_02_07 with 20598 reviewed entries). Precursor and fragment mass tolerances were set at 10 ppm and 0.02 Da respectively, and up to 2 missed cleavages were allowed. Oxidation (M) was set as variable modification, and Carbamidomethylation (C) as fixed modification. Peptides were filtered with a false discovery rate (FDR) at 1%, rank 1. Proteins were quantified with a minimum of 1 unique peptide based on the XIC (sum of the Extracted Ion Chromatogram). The quantification values were exported in Perseus^87^ for statistical analysis involving a log2 transform, imputation, normalization^87^.

### Proteome analysis (CRCM) Sample preparation (CRCM)

Briefly, 15µg of each protein extract (n=3, biological replicates) were loaded onto NuPAGE™ 4–12% Bis–Tris acrylamide gels according to the manufacturer’s instructions (Life Technologies). Electrophoresis was stopped as soon as the proteins had stacked in a single band. Protein-containing bands were stained with Imperial Blue (Pierce), excised from the gel, and digested with high sequencing grade trypsin (Promega) prior to mass spectrometry analysis. Briefly, gel pieces were washed and destained through several washed with 100mM NH4HCO3. Destained gel pieces were shrunk with 100 mM NH4HCO3 in 50% acetonitrile and dried at room temperature. Protein-containing del pieces were then reduced with 10mM DTT in 25 mM NH4HCO3 (pH 8.0) for 45 min at 56°C. This solution was replaced by 55 mM iodoacetamide in 25 mM NH4HCO3 (pH 8.0), and the gel pieces were incubated for 30 min at room temperature in the dark. They were then washed twice with 25 mM NH4HCO3 and finally shrunk by incubation for 5 min with 25 mM NH4HCO3 in 50% acetonitrile. The resulting alkylated gel pieces were dried at room temperature. The dried gel pieces were rehydrated by incubation in 25 mM ammonium bicarbonate (pH 8.0) supplemented with 12.5 ng/µl trypsin (Promega) for 1h at 4°C and then incubated overnight at 37°C. Peptides were recovered by collecting the initial digestion solution and performing two successive extractions: first with 5% formic acid and then with 5% formic acid in 60% acetonitrile. The pooled extracts were dried in a centrifugal vacuum system.

### Peptide separation and mass spectrometry analysis (CRCM)

Samples were analyzed using a label free quantification (LFQ) strategy in data independent acquisition mode (DIA) mode. LC–MS/MS analysis were performed using an Orbitrap Astral mass spectrometer (Thermo Fisher Scientific) coupled to an EVOSEP ENO Chromatographic system. The vacuum-dried peptide samples were resuspended in 0.1% of formic acid, and 1µg of peptides was subsequently loaded onto Evotip Pure™ tips according to manufacturer’s instructions. Peptides were separated using an Evosep performance column (Evosep, Cat. No. EV1137, 1.5µm beads, 150 μm × 15 cm) with the 30 Samples per day preprogrammed gradient. The column temperature was maintained at 40°C using an Evosep Pod for Thermo Easy-Spray source. DIA data were acquired on the Orbitrap Astral Mass Spectrometer using the following parameters. MS1 spectra were acquired in the Orbitrap analyser at a resolution of 240,000 over a mass range of 380-980 m/z, with a scan cycle time of 0.6 s. The MS1 normalized AGC target was set to 500%, with a maximum injection time of 5 ms. DIA MS2 spectra were acquired in the Astral analyzer over a mass range of 380-980 m/z, with a normalized AGC target of 500%, a maximum injection time of 3 ms, and an HCD collision energy setting of 25%. Window placement optimization was enabled. The isolation window was set to 2 Th without window overlap, and the fragment scan range was set to 150-2000 m/z.

### Data processing protocol and bioinformatics analysis (CRCM)

Relative intensity-based label-free quantification (LFQ) was processed using the DIA-NN 2.2 algorithm^88^. Raw files were searched against the Human database extracted from UniProt on the 3rd of March 2026 and containing 20 431 entries with the addition of a protein contaminant blank^89^ (Ref 3 Frankenfield et al., 2022). The following parameters were used for searches: (i) trypsin allowing cleavage before proline; (ii) one missed cleavage was allowed; (iii) cysteine carbamidomethylation (+57.02146) as a fixed modification and methionine oxidation (+15.99491) and N-terminal acetylation (+42.0106) as variable modifications; (iv) a maximum of one variable modification per peptide allowed; and (v) minimum peptide length was 7 amino acids and a maximum of 30 amino acids. The match between runs option was enabled. The precursor false discovery was set to 1%. DIA-NN parameters were set on Single-pass mode for the Neural Network classifier, Robust LC High precision for quantification strategy and RT-dependent mode for Cross-run normalization. The library was generated using a Smart profiling setup. The main output file from DIA-NN was further filtered at 1% FDR and LFQ intensity was calculated using our DIAgui package^90^ at 1% q-value.

Quantification values were further treated with Perseus^87^ for filtering, imputation and statistical tests, using mostly default settings. For filtering, the minimum percentage of values was set at 70% in at least one group. The FDR was set at 0.05 using Benjamini-Hochberg correction. Multiple sample testing (ANOVA) was run on significant results using default values followed by a post-hoc Tuckey HSD with FDR of 0.05.

### NleA purification

His-mCherry-Avi-NleA and His-mCherry-Avi-NleAΔPBM synthetic genes were purchased from Twist Bioscience, in an *E. coli* optimized vector.

Transformation was performed by mixing 1µl of plasmid with 25 µl of competent cells, incubated on ice for 20min then heat shocked at 42°C for 45s, recovered in SOC media at 37°C for 1h, then used to prepare glycerol stocks (50%) that were stored at -80°C. Precultures from glycerol stocks were grown at 37 °C overnight in 50ml LB with Kanamycin and Chloramphenicol and 5x10ml were used to inoculate 5x400ml 2YT media (Sigma), grown at 37°C until OD ∼0.75 then induced with 1mM IPTG and cultured overnight at 25°C. Cells were recovered by centrifuging at 4,000g for 10min, resuspending the pellet in lysis buffer (Tris 50mM, NaCl 300mM, pH 8, 0,5 mg/mL lysozyme) and stored at -20°C.

Purification of NleA and NleAΔPBM was performed as for individual PDZ domains (described above) except only Ni^2+^ affinity was used (5ml Ni FF crude, on Akta Express). Elution and wash fractions were pooled, concentrated to 5ml total volume and injected into a gel filtration TSK3000 column, after which monomer fractions for NleA and NleAΔPBM were separately pooled and stored as 4x1ml at 3.5mg/ml and 14x1ml at 2.2 mg/ml, respectively.

### NleA biotinylation and native Holdup experiment

Aliquots of NleA and NleAΔPBM, were recovered from the -20°C freezer and biotinylated for native Holdup experiments. The biotinylation reaction was carried out at room temperature for 1h under agitation, by diluting both NleA and NleAΔPBM to 24µM in a solution containing 1.2 µM BirA (1:20), 5mM ATP, 36 µM biotin (1.5:1) and 10mM MgSO4.

Afterwards, samples were desalted with Nap5 columns (500ul sample, Cytiva) to remove excess biotin and then incubated with 25ul streptavidin resin for 2h at 4°C in 0.5ml protein LoBind tubes (3 tubes per protein).

After incubation, the protein-bound resin was washed 3x in 400µl native Holdup buffer (50 mM TRIS pH 7.5, 300 mM NaCl, 1 mM TCEP), then all supernatant was carefully removed and a singlicate of each protein was incubated with 100µl of cell extract from Caco2, HEK293 and polarized Caco2 cells, for 2h at 4C. After incubation, 25µl were collected for MS analysis as described above.

### Fragmentomic Holdup data analysis

Holdup assay data was analyzed as in^41^, using the same equations for binding intensity (BI) calculations, determination of dissociation constants (Kd) and Gibb’s free energies (ΔG and ΔΔG) The binding intensity threshold was set at 2 standard deviations of the mean binding intensity for all biotin controls (0.026). BI values were corrected with a normalization factor to account for PDZ integrity, calculated from capillary electrophoresis analysis of purified PDZ domains.

### Native Holdup data analysis

Native Holdup data was analyzed as in^55^. First, the standard deviation (σ) of log2(fold-change) values was calculated and a hard threshold was placed at -1σ. For *P values*, the statistical significance threshold was set at 0.05 (-log10(*P value)* > 1.3). Then, two hyperbolic threshold curves were plotted at two different curvatures: c=1 (as in^55^) and c=0.1. For details about calculations see Zambo et. al^55^. The more relaxed threshold at c=0.1 was set based on the agreement between negative hits for both fragmentomic and native Holdup experiments. Affinities were calculated by assuming prey concentrations of 100nM and effective bait concentrations of 10µM using the same equations as in^55^.

An orthogonal annotation of all depleted proteins was implemented to determine if some of them might have been “dragged” by direct interactors of PBMs fixed on the resin. To this end, we downloaded the full set of human PPIs from IntAct^91^ (downloaded on August 16,2024), filtered it to keep only interactions annotated as “association”, “physical association” or “direct interaction” and created interaction networks from these for depleted proteins in each run. If a depleted protein is found within a connected component that includes a PDZ containing protein that was also depleted, it was marked as an indirect interactor.

Similarly, to determine if any of the motifs overlapping with C-terminal PBMs could be driving certain interactions with depleted proteins, we annotated all depleted proteins with their corresponding PFAM domains and checked if these corresponded to targets of the identified motifs. When true, these depleted proteins were annotated as “ELM other”.

### *In cellula* experiments

#### Cloning

The plasmid pcDNA5-PDZK1-HA was obtained from MGC BioCampus (Montpellier, France). Plasmids pTWIST-NleA and pTWIST-NleAΔPBM were obtained from XX. NleA and NleAdPBM were amplified by PCR using Phusion high-fidelity polymerase (Thermo Fisher Scientific) and primers NleA-EcoRI-Fw (CTCGAATTCTAATATACAGCCCATTGTTACC) And NleA-BamHI-Rv (GTTGGATCCTCACACCCGTGTTTCCTG) Or NleAdPBM-BamHI-Rv (GTTGGATCCTCAATCAACATAATTAGATGGG), respectively. PCR products were purified and digested with EcoRI and BamHI (New England Biolabs). Digested PCR products were ligated into pLVX-GFP-N2 (Burette et al., 2020) to generate pLVX-GFP-NleA and pLVX-GFP-NleAdPBM. Plasmid sequences were verified by DNA sequencing (Eurofins).

#### Cell culture and transfection

U2OS (ATCC) cells were routinely maintained in DMEM containing 10% fetal calf serum (FCS) in a humidified atmosphere of 5% CO2 at 37° C.

Cells were grown to 70% confluence in 96 well plates (black µclear, Greiner Bio One) and transfected with JetPEI cationic polymer transfection reagent (Polyplus Transfection) diluted in 150 mM NaCl, according to the manufacturer’s recommendations.

#### Immunofluorescence staining

Cells were fixed in 4% (wt/vol) paraformaldehyde in phosphate-buffered saline (PBS) pH 7.4 solution at room temperature for 20 min. Samples were then rinsed with PBS solution and incubated in blocking solution (0.5% bovine serum albumin [BSA], 50 mM NH4Cl in PBS solution, pH 7.4). Cells were then incubated with rat anti-HA primary antibody (Cell Signalling) diluted in blocking solution for 1 h at room temperature in a humidified atmosphere, washed five times in PBS solution, and further incubated for 1 h with anti-rat secondary antibody coupled to Alexa Fluor 555 (Thermo Fisher Scientific) diluted in blocking solution complemented with Hoechst 33258 (Sigma) for DNA staining. Cells were washed as previously described and maintained in PBS for image acquisition.

#### Image acquisition and analysis

Images were acquired using an automated high-resolution confocal microscope (Opera Phenix^TM^ high-content imaging systems, Revvity) with a 20X Air NA.08 objective. Images were acquired in EGFP, Alexa 568 and Hoechst channels every 2 µm across an 8 µm section in 14 positions per well. For each position, stack images were then processed for maximum intensity projection.

The Harmony software (v5.3, Revvity) was used for object identification and signal overlap analysis.

#### Statistical analysis

Statistical analysis was performed using Prism (GraphPad).

#### T3SS and T4SS Sequences

A list of pathogen T3SS and T4SS secreted effector sequences had been previously downloaded from BastionHub^92^ (versions from July 07, 2020) and complemented with manual annotation of effectors from the literature. This list of sequences was used as input for the selection of C-terminal PBMs. Briefly, C-terminal PBMs were selected by matching the PDZ ligand regular expressions from the ELM database^40^ for LIG_PDZ_Class_1, LIG_PDZ_Class_2, LIG_PDZ_Class_3 and LIG_PDZ_Wminus1_1, with regular expressions `…[ST].[ACVILF]$`, `…[VLIFY].[ACVILF]$`, `… <ED>.[ACVILF]$` and `.w[ACGILV]$`, respectively. Fasta files were opened with the SeqIO module from the Bio library in Python and regular expressions were matched using the re library. The ELM database reports regular expressions comprising six positions for PBMs in classes 1-3. The canonical human proteome contains 3749 distinct six amino acid PBMs in 4115 proteins. Out of the ∼200 selected bacterial PBMs, only 5 of them are an exact match to a human PBM over six positions (supfig.1, c): the VQDTRL C-terminal of effector Map from pathogenic *E. coli* and homologs, matching the last six amino acid positions of the human cystic-fibrosis transmembrane regulator protein (CFTR). Although positions 0 to -3 from the C-terminal are the most important for PDZ-PBM binding, sequence context beyond the last four positions contributes to specificity and binding affinity^37,93^. As such, our selection of 10 amino acid long PBMs allows probing binding specificities beyond the definitions in ELM, potentially revealing evolutionarily tailored targeting of PDZ domains by bacterial effectors.

### Node degree comparison

To determine if PDZ domains with many interactions (network hubs) in the human PDZ-PBM interactome reported in Profaff^41^ (http://profaff.fr/index.php) tend to be hubs in the bacterial PDZ-PBM network as well, we calculated the normalized node degree for both datasets as:

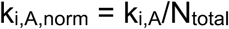

where ki,A,norm is the normalized degree of node I in network A, ki,A is the degree of node I (number of interactions) in network A and Ntotal is the total number of PBMs tested in the database and calculated a simple bias metric as:

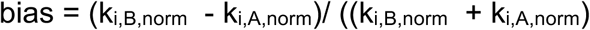

where ki,B,norm and ki,A,norm are the number of interactions of node i with PBMs in our dataset (B) and in Profaff^41^ (A).

### Interaction similarity clusters

A clustering analysis for both PDZ and PBM was performed based on interaction similarity matrices between affinity vectors we calculated previously (PDZ vs PDZ, PBM vs PBM). Briefly, we represented the merged interaction data from both screens as a network and used the affinity weighted adjacency matrix to calculate a modified Dice index (see methods) for all possible PBM affinity vector pairs (624^2) and clustered the resulting dissimilarity vectors together.

We performed a centroid based (Kmeans) and a hierarchical clustering of these matrices to create clusters of PDZ domains and PBMs with similar interactions profiles. Kmeans allows for a broader grouping of PBMs based on interaction similarity, while hierarchical clustering can discern if certain residues drive specificity for highly similar PBMs. For the Kmeans analysis, the number of clusters was selected using the “Elbow method”. First, we ran the Kmeans algorithm for a range of values from 1 to 50, calculating the inertia (within cluster sum of squares) for each one and plotting the resulting inertia curve. A line connecting the first and last points in the range was traced and we calculated the orthogonal projection of each point in the inertia curve onto said line. The point with the orthogonal projection maximizing the distance between the curve and the line was chosen as the number of clusters. For Kmeans we used the Python library Sklearn’s (v 1.6.1) KMeans module. For hierarchical clusters we used Seaborn’s (v 0.13.2) clustermap using a Euclidean metric and average linkage.

### Interaction dissimilarity calculation

From the PDZ-PBM interaction network we extracted individual interaction affinity vectors for all PDZ domains and PBMs and computed a modified Dice index by rewriting the equations from^94^ in vector form and computing the individual dissimilarity measures for each pair of PDZ-PDZ or PBM-PBM affinity vectors. The calculation is divided into terms accounting for common interactors (attraction term) and interactors unique to either member of the pair (repulsive term):

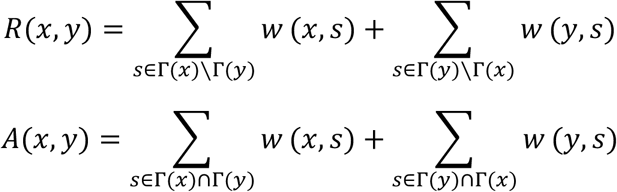

where **x, y** and **s** are nodes (either PDZ or PBM), and:

*R*(*x*, *y*) is the repulsive part,

*A*(*x*, *y*) is the attractive part,

*w*(*x*, *s*) are the weights of the interaction between x and s,

*w*(*y*, *s*) are the weights of the interaction between y and s,

Γ(*x*) and Γ(*y*) are the neighborhoods of x and y, respectively.

we can rewrite these as:

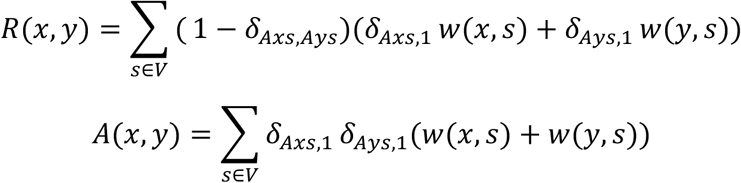

where x and y are nodes, and:

*A_xs_* is an entry between nodes x and s in the adjacency matrix **A**,

*δ_x_*_,*y*_ is the Kronecker delta (= 1 if *x* = *y*, 0 otherwise). and finally cast these into vector form:

First, we define the Hadamard product:

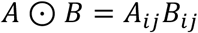

where A and B are matrices.

For vectors *x⃗* and *y⃗* this is the elementwise product that keeps all common interactions:

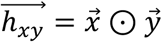

We consider *x⃗* and *y⃗* to be unweighted row vectors from the adjacency matrix A. In other words, *x⃗*, *y⃗* ∈ [0,1]*^n^*.

Now we can express *A*(*x*, *y*) and *R*(*x*, *y*) as:

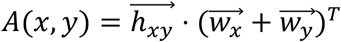

where 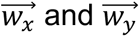 are the weights associated to *x⃗* and *y⃗*, respectively.

For *R*(*x*, *y*), we keep only elements that are 0 in one and 1 in the other. We can do this by summing both binary vectors and keeping only elements equal to 1 (the sum ∈ [0,1,2]*^n^*).

Let *z⃗* = (*x⃗* + *y⃗*) mod2, then:

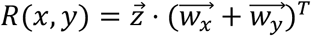

Finally, we calculate dissimilarity as:

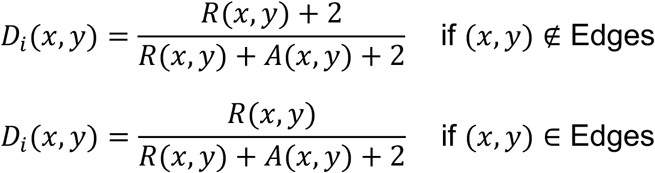

### Individual domain targeting

A targeting analysis based on affinity distributions was performed to compare bacterial pathogenic and commensal effector PBMs to previously published PDZ-PBM data, between human PDZ domains and human and viral PBMs. Among the 253 PDZ domains we measured interaction affinities for, 128 are also present in the Profaff database^41^. We grouped PBMs into bacterial commensal (resident gut bacteria), bacterial pathogen, human and viral groups, and compared the distributions of their interaction affinities for common PDZ domain targets (Fig.2). Kruskal-Wallis, Dunn’s correction, P-value<=0.05) (Fig 2.).

Overall affinity distributions for PBMs from each group (bacterial commensal, bacterial pathogenic, human and viral) with each human PDZ domain were compared using the non-parametric Kruskal-Wallis test with Dunn’s correction for multiple testing, setting a *P value* <=0.05 for significance, to assess preferential targeting of human PDZ domains for all tested PBMs and potential PPI competition hotspots. Data were compared for any given PDZ domain if at least five data points from two groups were above the selected affinity threshold. Of the initial 128 PDZ in common in both studies, 65 show no significant differences, 23 were not comparable due to insufficient data and the remaining 40 showed statistically significant differences between at least two groups.

### Gene set enrichment analysis

Gene set enrichment analysis (GSEA)^95^ was performed to compare if for certain groupings of the HU data some PDZ domains were overrepresented at the top or bottom of the affinity list. First, we ordered the interaction data by decreasing affinity then grouped them according to interaction similarity clusters and used the GSEApy^96^ Python package (v1.1.0) to calculate enrichments using the “prerank” modality with 1000 permutations. The false discovery rate q-value (FDR q-val) threshold was set at 0.15.

### PSSM

We used pLogo (v1.2.0)^53^ to calculate a position weighted scoring matrix (PWSM) for each interaction similarity cluster, by taking all PBMs interacting with PDZ members of each cluster and setting an affinity threshold at pkd>4 (Kd <100μM) . pLogo corrects for frequency biases in amino acid composition at each position, allowing for the generation of frequency corrected sequence logos. The background for calculation of the PWSM was the full set of tested effectors.

### Functional enrichment

Functional enrichment was performed using gProfiler^97^ on all interaction similarity clusters.

For PDZ in the Profaff database^41^ and in our screen, interaction data from both datasets was merged. Prior to the analysis, all human proteins with PDZ domains or C-terminal PBMs were annotated with tissue protein expression data from the Human Protein Atlas^47^ (downloaded in May 13, 2023) and bacterial effector proteins were annotated according to tissue tropism. The interaction dataset was filtered to include gastrointestinal tract and lung tissues and split accordingly.

Functional enrichment was performed by grouping all human PBMs in the interaction similarity clusters along with their target PDZ domains for all interactions with affinities above a pkd of 4 (<100µM) for both GI tract and lung datasets. The choice of affinity threshold is based on the interaction assay recovery rates for the HU assay (supplementary figure 3,). The enrichment was performed against the set of PDZ and PBM interactors present in Profaff^41^ (FDR, *P value* < 0.05), against a set containing all 152 human PDZ containing proteins and all ∼4000 canonical human PBMs (FDR, *P value* < 10^-3^ and 10^-2^) and against the full human proteome (FDR, *P value* < 10^-5^).

For semantic similarity clustering and visualization of enrichment terms, we used the Revigo^54^ R library, with a redundancy threshold at 0.7 and the Bioconductor “org.Hs.eg.db” database.

### Python library versions

Unless otherwise indicated, the following versions of Python libraries were used for analysis: Scipy v1.13.1, numpy v1.26.4, pandas v2.2.3, matplotlib v3.8.3, sklearn v1.6.1, Seaborn v0.13.2, Bio v1.85, re v2.2.1

## Supporting information

Supplemental Fernandezmacgregor

## Author contributions

Project conception: CB, AZ, RV; PDZ production: J.F.M, L.I., J.M.D, F.B., R.V.; 14-3-3 production: J.F.M, L.I., R.V.; Holdup assay: J.F.M, L.I., R.V.; Peptide synthesis: S.O.; native Holdup: J.F.M, E.M., B.M., G.T., V.K., C.G.; Mass spectrometry: B.M., S.A, L.C, J.-P.B; T3SS and T4SS selection: J.F.M, A.Z.; Intact interaction interface annotation: L.B.; Holdup analyses and native Holdup analyses: J.F.M.; *In cellula* experiments: E.Ma., A.B., V.M.; manuscript writing: J.F.M., R.V., A.Z., C.B.; Funding acquisition: R.V., C.B., A.Z., G.T., J.F.M

## Acknowledgements

The project leading to this publication has received funding from France 2030, the French Government program managed by the French National Research Agency (ANR-16-CONV-0001), from the French National Research Agency (ANR-22-CE44-0018), from Excellence Initiative of Aix-Marseille University – AMIDEX (AMX-21-PEP-043), and also from the JPI HDHL-INTIMIC action co-funded by the Agence Nationale de la Recherche (ANR-17-HDIM-0001). J.F.M. was funded by Consejo Nacional de Humanidades Ciencias y Tecnologias (CONAHCYT) Becas al Extranjero Convenios GOBIERNO FRANCES 2021 -1 grant 795494, and received support from the ‘Espoirs de la Recherche’ program managed by the French Fondation pour la Recherche Medicale (FRM, FDT202404018637). The AFMB was supported by the French Infrastructure for Integrated Structural Biology (FRISBI) ANR-10-INSB-05-01; Marseille Proteomics (marseille-proteomique.univ-amu.fr) was supported by IBISA (Infrastructures Biologie Santé et Agronomie), Plateforme Technologique Aix-Marseille, the Cancéropôle PACA, the Provence-Alpes-Côte d’Azur Region, the Institut Paoli-Calmettes, the French National Cancer Institute (INCa), the European Regional Development Fund (FEDER) and the French Plan Cancer. This work was also supported by the National Centre for Scientific Research (CNRS) and Aix-Marseille University. Diala Kantar and Etienne Coyaud (PRISM, Lille) kindly provided HEK293 cell extracts for native Holdup at IGBMC; Claire Debarnot and Hussein Hamade kindly provided HEK293 and Caco2 and differentiated Caco2 cells for extract preparation for native Holdup at CRCM.

## Data availability

The Astral mass spectrometry proteomics data have been deposited to the ProteomeXchange Consortium via the PRIDE partner repository under the dataset identifiers PXD0ZZZZZ, PXD0XXXXX, and PXD0YYYYY.

