## Supplemental Fernandezmacgregor for "A quantitative interactome map reveals affinity tailored hijacking of human cellular functions by resident and pathogenic bacteria"

### Supplementary Information

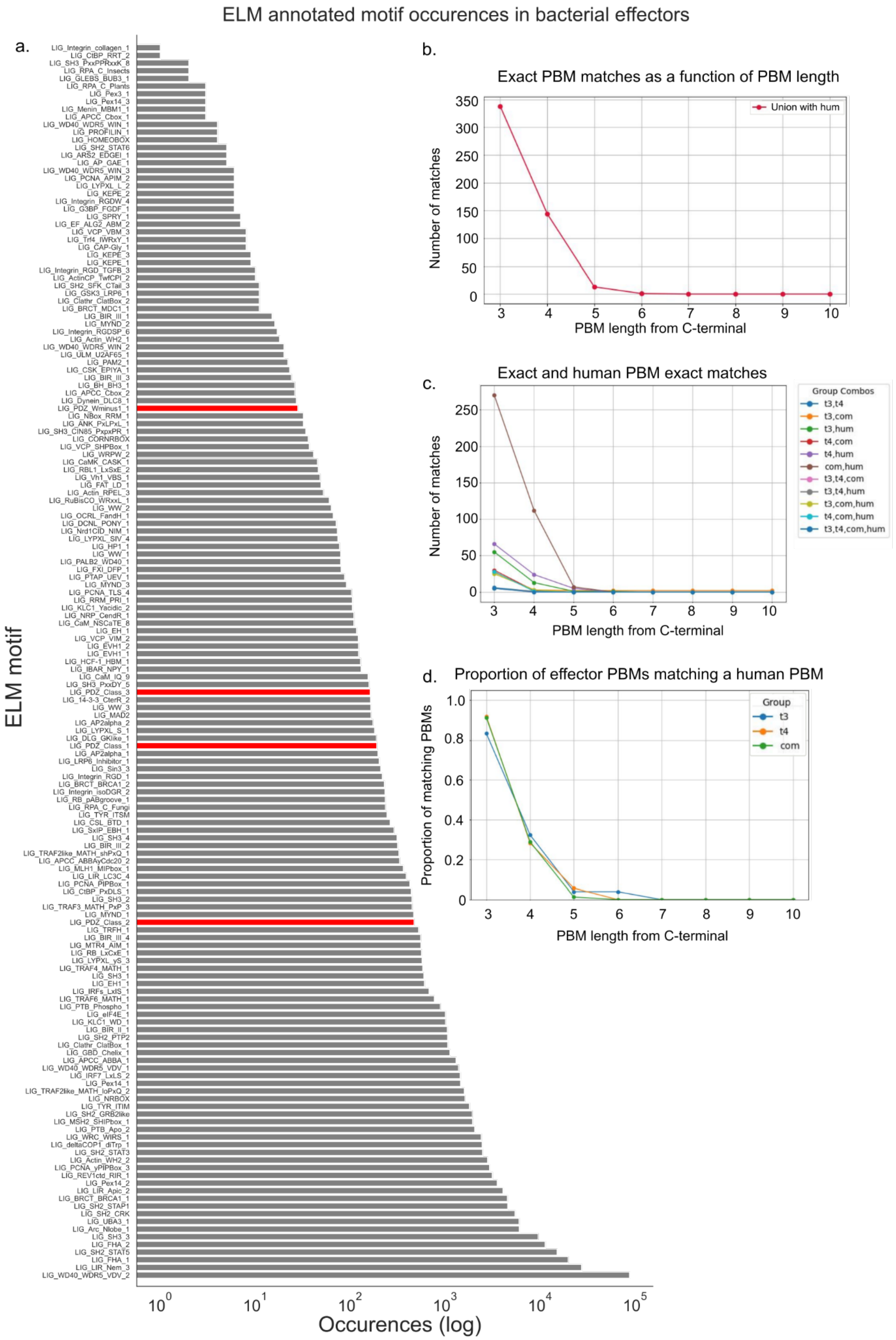

**Supplementary Figure 1 | Bacterial effectors SLiMs description and PBM comparison to human PBMs. a,** ELM motif counts for all effectors in the initial set (~4900) with PDZ-binding classes shown in red. **b,** Counts of PBM exact sequence matches between bacterial and human PBM as a function of position from the C-terminal (the high number of overlaps up to positions 3 and 4 are a result of the imposed regular expression profiles). **c,** number of exact sequence matches for human and bacterial PBMs, separated by all possible group combinations with commensals in a single group and pathogen effectors separated by secretion system. **d,** Overall proportion of effectors with an exact match to a human PBM as a function of PBM length from the C-terminal.

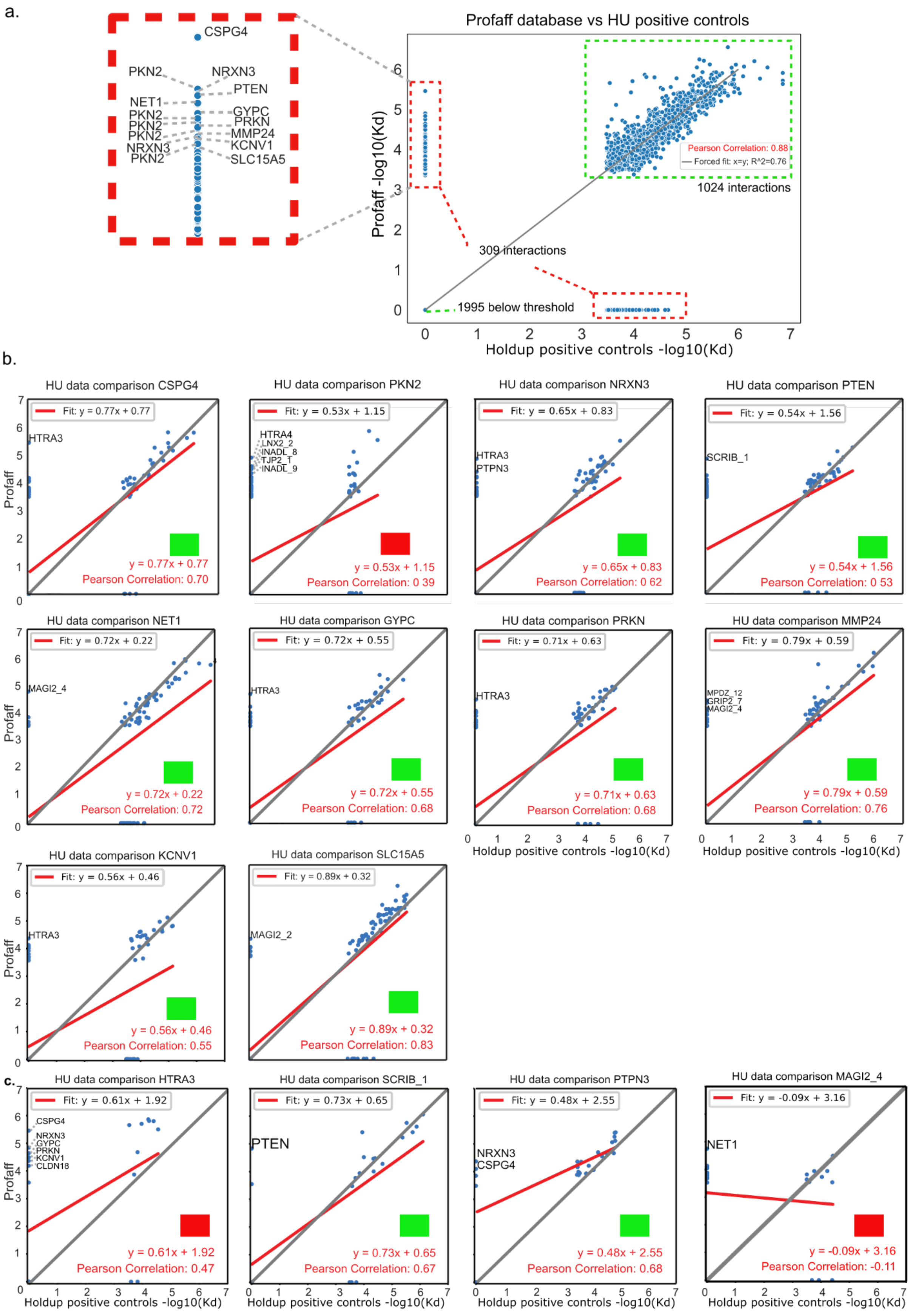


**Supplementary Figure 2 | Exploration of positive controls mismatch region. a, (right)** Scatter plot of full set of positive control PBMs vs the corresponding PBMs reported in the Profiling affinities database[1], with match and mismatch regions highlighted in green and red, respectively. Each point corresponds to the interaction of the same PBM in both datasets against a given PDZ domain. **(left)** Close-up of the higher end of the affinity range mismatch region with PBM names shown. Because the recall rate for the lower end of the affinity range varies between 50 and 80%, mismatches in this region are expected. Mismatches in the higher affinity range could indicate problems with control PBMs or the corresponding PDZ domains. **b, PBM** Scatter plot of the 10 individual PBM positive controls (out of 24 total) shown for the high affinity inset region (a, left) with individual points in the high affinity mismatch region annotated with the corresponding PDZ domain names, revealing a control with low quality fit highlighted by a red square (e.g. PKN2) and controls with good fits highlighted by a green square showing a few outliers. NOTE: x and y axes are the same for all plots. **c, PDZ** Scatter plots of individual PDZ domains corresponding to outliers identified for good quality positive controls, revealing good quality fits highlighted by a green square for SCRIB_1 and PTPN3 and poor fits highlighted in red indicating a problem with PDZ domains HTRA3 and MAGI2_4. Jointly, the HTRA3 and MAGI2_4 PDZ domains and the PKN2 PBM explain the majority of mismatches in the high affinity region.

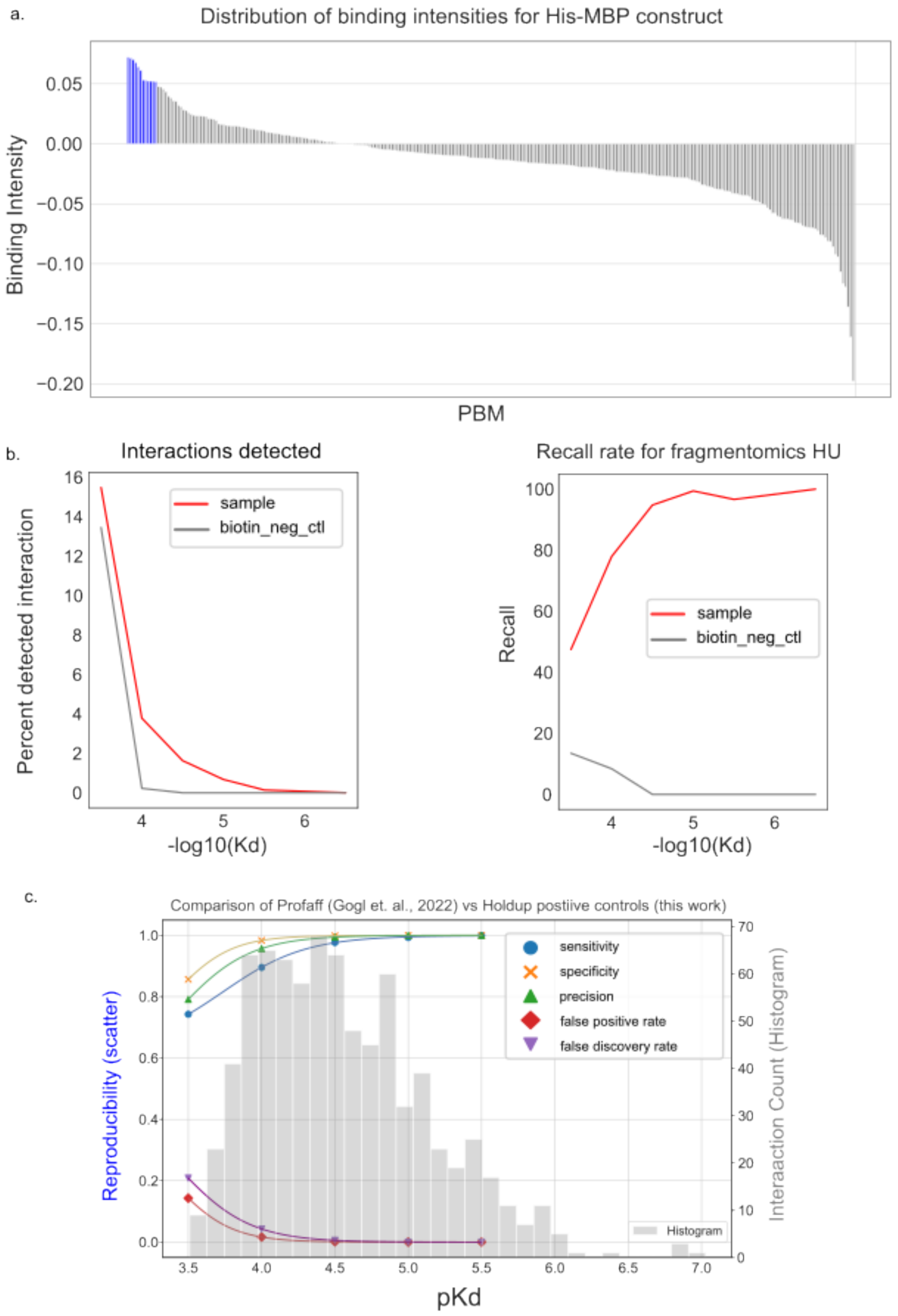


**Supplementary Figure 3 | HU controls. a,** HIS-MBP construct interactions against 289 PBMs and biotins, only 12/289 (~4%) points are barely above the median affinity threshold for all runs, indicating the HIS-MBP construct does not interact with PBMs or biotin controls. **b,** interaction curves as a function of affinity for both samples (red) and biotin negative controls (grey) (left) and the corresponding recall curves for all replicates (for samples a total of 41433 replicates were measured for 15907 binary PDZ -PBM interactions for 44 PDZ domains) (right). The false positive rate for the low affinity region of the graph is ~13% with an associated recovery rate of ~14% compared to ~16% interactions detected for samples with ~47% recovery. Above -log10(Kd) values of 4 (Kd < 100μM) biotin controls practically drop to zero while samples are reliably detected with recovery rates between 80-100%. **c,** Holdup assay sensitivity, precision, specificity, FDR and FPR rates for positive controls compared to Profaff database[1] (corresponds to scatter plot in main Fig.1). The increase in overall confidence for pKd>4 reinforces the notion of a grey zone for the lower end of the affinity spectrum and justifies performing affinity comparisons for interactions with pKd>4

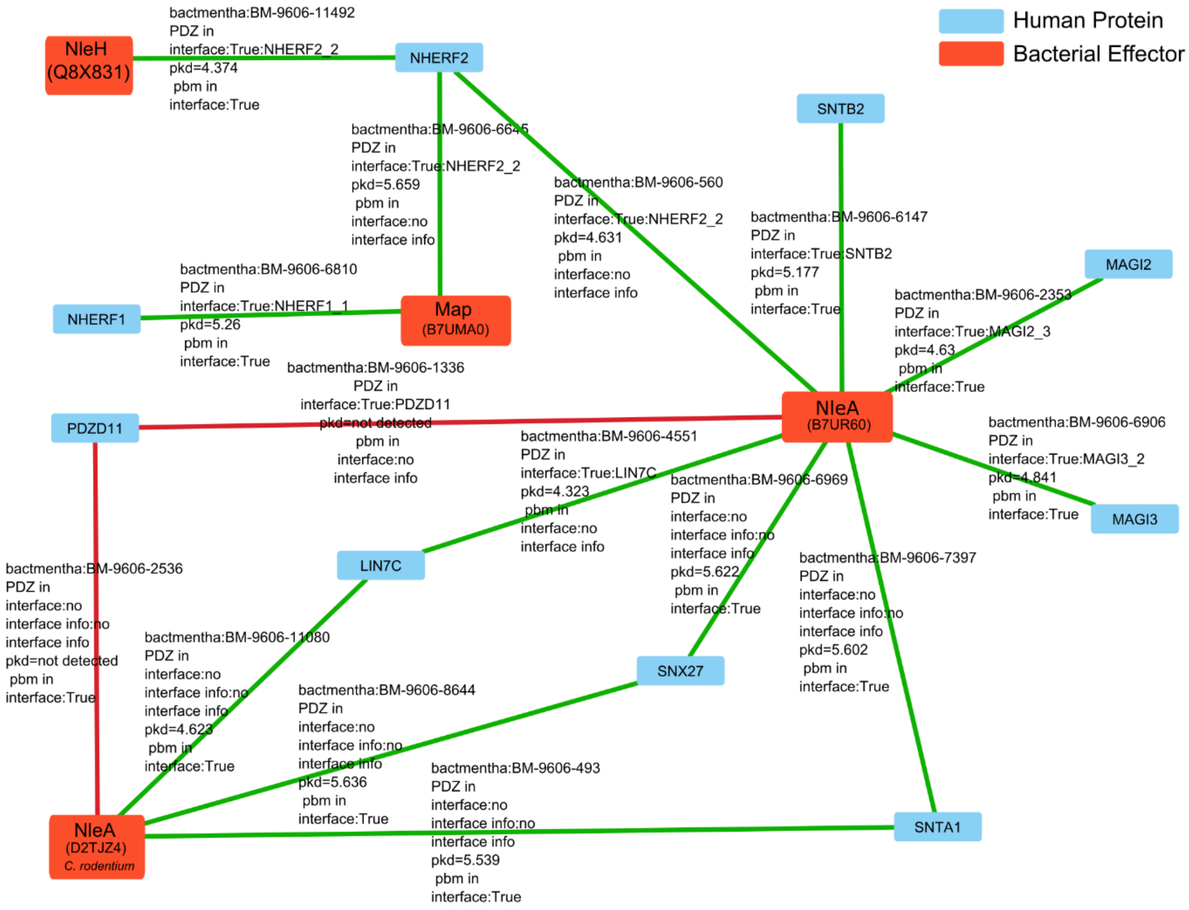


**Supplementary Figure 4 | Recovery of interfaces via HU for PPIs reported in Intact[2].** Validation of full-length PPIs reported in the literature involving either a human PDZ domain (blue square nodes) and/or a bacterial PBM (bright orange square nodes) at the interaction interface. Interactions (edges) for which a corresponding PDZ-PBM interaction was identified by Holdup are shown as green links(13/15 ~87%) and those for which no interaction was found are shown as red links.

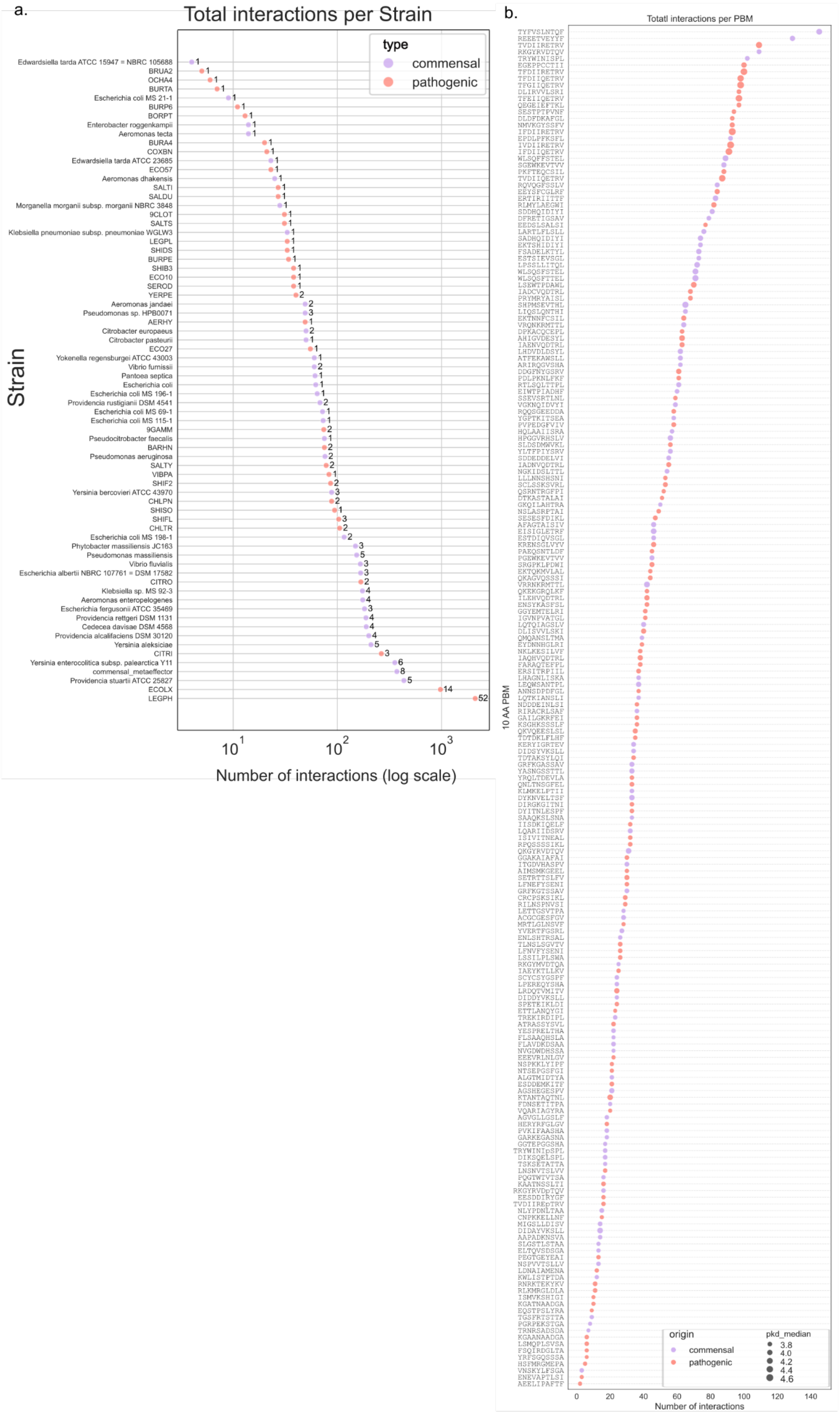

a.
Total interactions per Strain
type
commensal
pathogenic
Strain
Edwardsiella tarda ATCC 15947 = NBRC 105688
BRUA2
OCHA4
BURTA
Escherichia coli MS 21-1
BURP6
BORPT
Enterobacter roggenkampii
Aeromonas tecta
BURA4
COXBN
Edwardsiella tarda ATCC 23685
ECO57
Aeromonas dhakensis
SALTI
SALDU
Morganella morganii subsp. morganii NBRC 3848
9CLOT
SALTS
Klebsiella pneumoniae subsp. pneumoniae WGLW3
LEGPL
SHIDS
BURPE
SHIB3
ECO10
SEROD
YERPE
Aeromonas jandaei
Pseudomonas sp. HPB0071
AERHY
Citrobacter europaeus
Citrobacter pasteurii
ECO27
Yokenella regensburgei ATCC 43003
Vibrio furnissii
Pantoea septica
Escherichia coli
Escherichia coli MS 196-1
Providencia rustigianii DSM 4541
Escherichia coli MS 69-1
Escherichia coli MS 115-1
9GAMM
Pseudocitrobacter faecalis
BARHN
Pseudomonas aeruginosa
SALTY
VIBPA
SHIF2
Yersinia bercovieri ATCC 43970
CHLPN
SHISO
SHIFL
CHLTR
Escherichia coli MS 198-1
Phytobacter massiliensis JC163
Pseudomonas massiliensis
Vibrio fluvialis
Escherichia albertii NBRC 107761 = DSM 17582
CITRO
Klebsiella sp. MS 92-3
Aeromonas enteropelogenes
Escherichia fergusonii ATCC 35469
Providencia rettgeri DSM 1131
Cedecea davisae DSM 4568
Providencia alcalifaciens DSM 30120
Yersinia aleksiciae
CITRI
Yersinia enterocolitica subsp. palearctica Y11
commensal_metaeffector
Providencia stuartii ATCC 25827
ECOLX
LEGPH
$10^1$
$10^2$
$10^3$
Number of interactions (log scale)
b.
Totatl interactions per PBM
10 AA PBM
origin
commensal
pathogenic
pkd_median
3.8
4.0
4.2
4.4
4.6
0
20
40
60
80
100
120
140
Number of interactions

**Supplemental Figure 5 | Number of interactions for bacterial effector PBMs. a,** Total number of interactions per bacterial strain (x-axis, log-scale) for resident microbiota (purple dots) and pathogen strains (salmon dots) with the number of distinct 10 amino acid PBMs shown on the right of each dot (for instance we tested 52 different PBMs *Legionella Pneumophila*), **b,** total number of interactions per PBM, with PBM sequence shown on the y-axis.

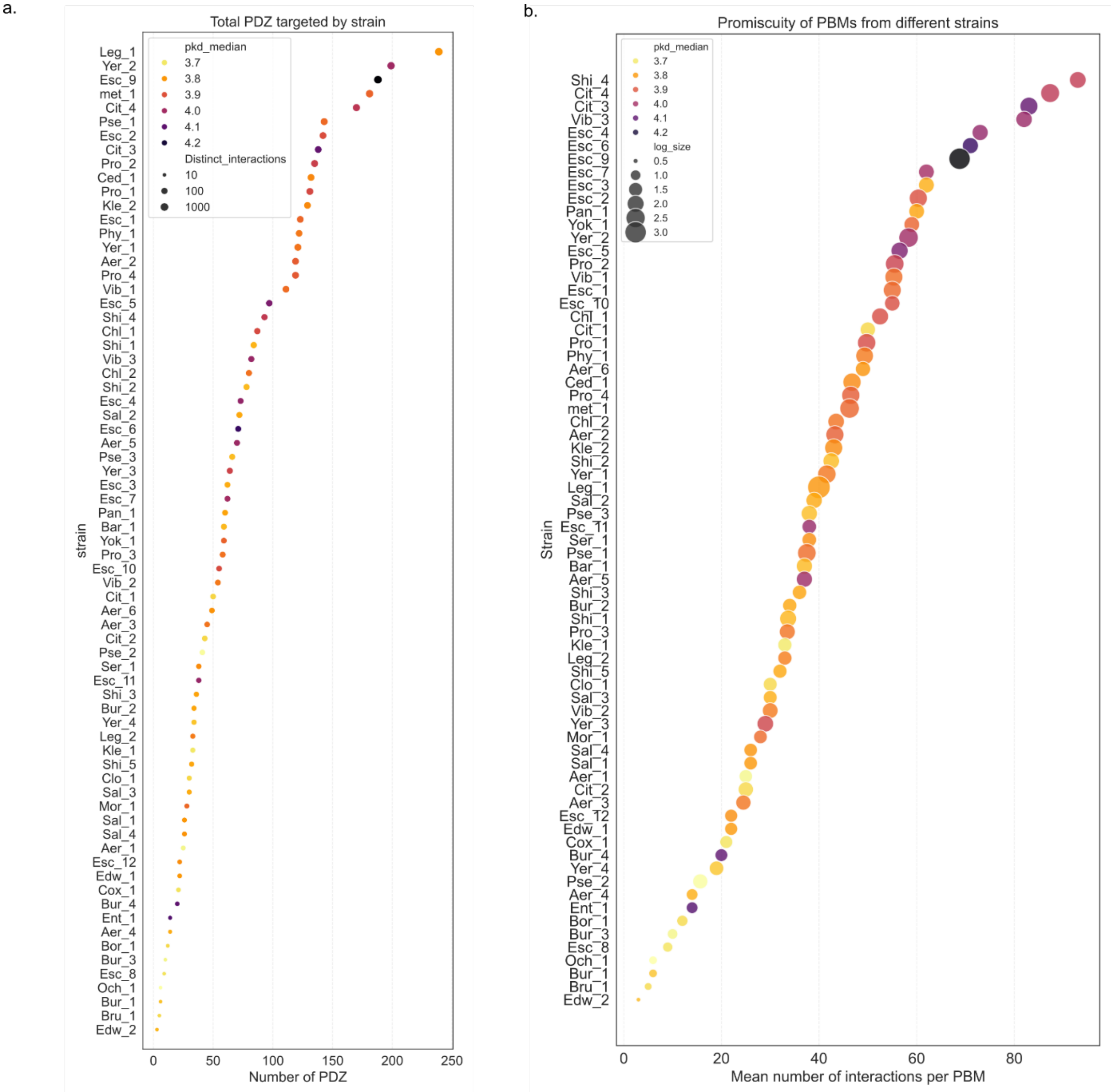


**Supplementary Figure 6 | Per strain coverage of human PDZome and PBM binding promiscuity for all affinities. a,** Experimental coverage of the full human PDZome by different strains for all quantified affinities (0.25-330 μM) and **b,** average number of significant interactions per PBM for each strain with dots colored by median pkd and sized by log10 of the total number of interactions for the strain.

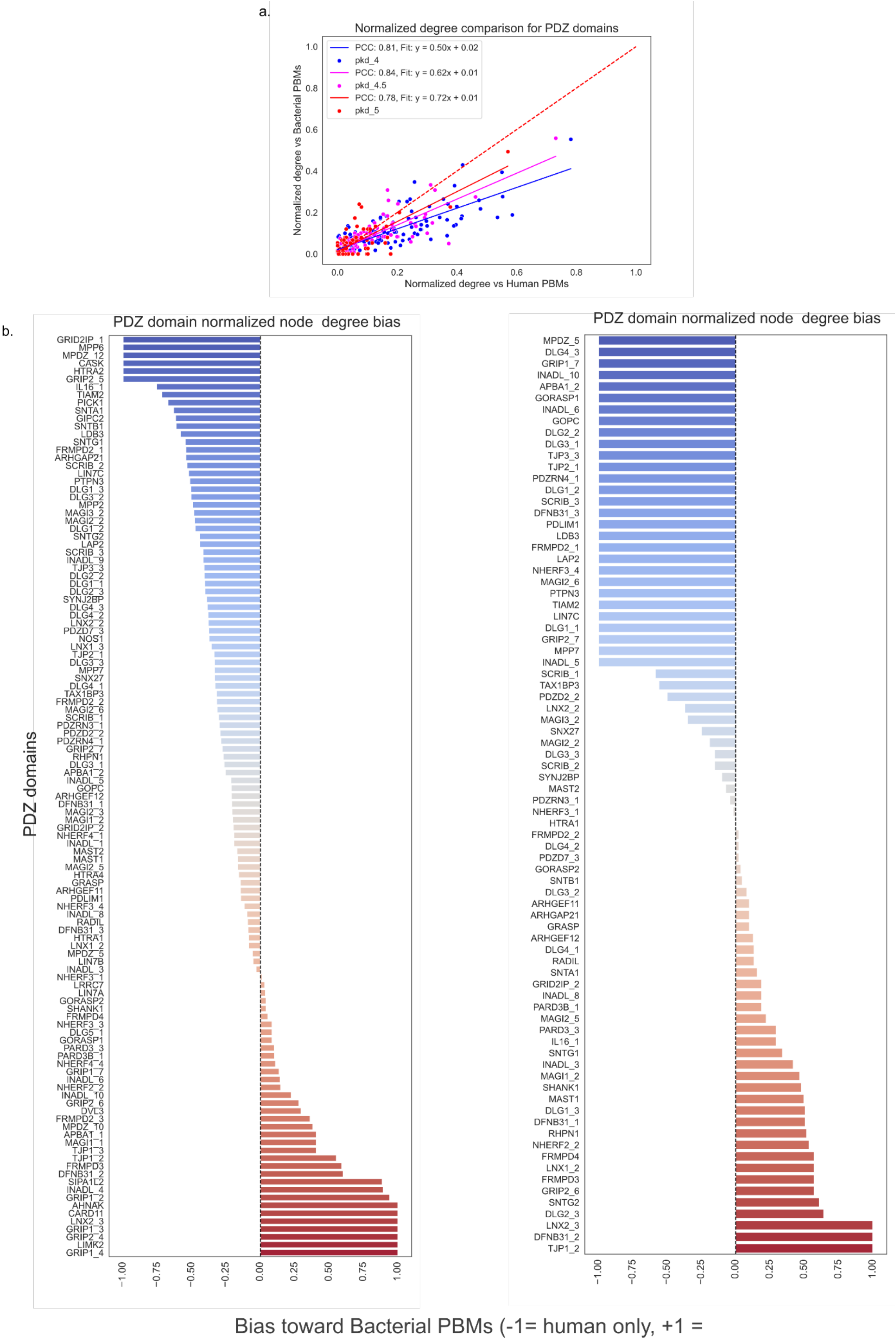


**Supplementary Figure 7 | Number of interactions for each PBM. a** normalized number of interactions for each PDZ domain (node degree in the PPI network) by total number of PBMs tested in our dataset (y-axis) and in Profaff (x-axis) for three different pkd thresholds. The dotted line corresponds to x=y and individual fits for each pkd threshold are shown along with PCC values. For all pkd thresholds, a linear correlation is observed between normalized node degrees indicating hubs in the human PDZ-PBM network are also hubs in the PDZ-bacterial PBM network, with a higher overall number of binders for each PDZ with human PBMs, compared to bacterial ones. **b,** targeting bias for PDZ domains by PBMs for either dataset for all interactions with affinities pkd >4 (left) and pkd > 5 (right). The pKd>5 plot highlights strong interactions in which a bias exists towards bacterial or human PBMs. Any PDZ domain for which the bias >-1 could have biological relevance.

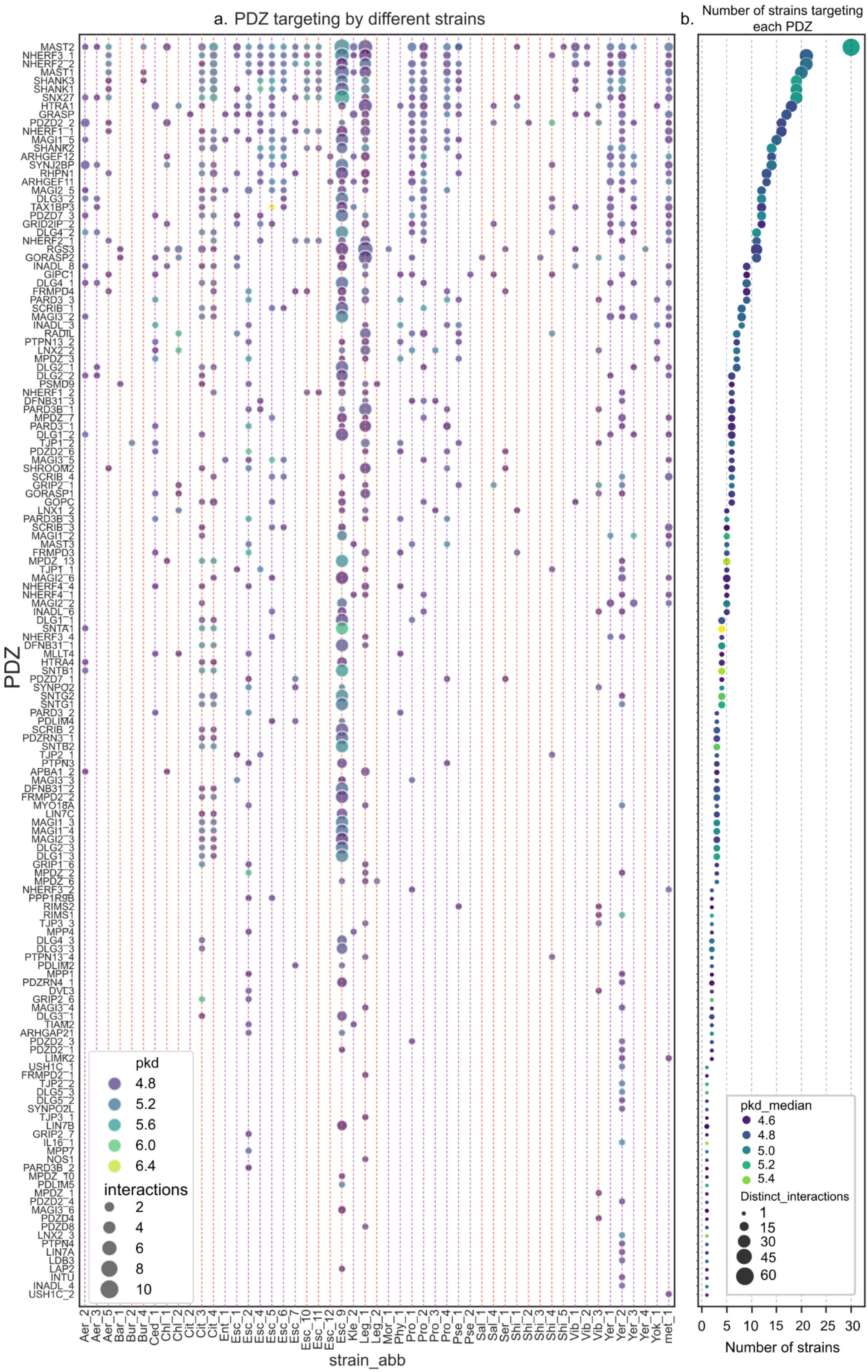
a. PDZ targeting by different strains
b. Number of strains targeting each PDZ
PDZ
MAST2
NHERF3_1
NHERF2_2
MAST1
SHANK3
SHANK1
SNX27
HTRA1
GRASP
PDZD2_2
NHERF1_1
MAGI1_5
SHANK2
ARHGEF12
SYNJ2BP
RHPN1
ARHGEF11
MAGI2_5
DLG3_2
TAX1BP3
PDZD7_3
GRID2IP_2
DLG4_2
NHERF2_1
RGS3
GORASP2
INADL_8
GIPC1
DLG4_1
FRMPD4
PARD3_3
SCRIB_1
MAGI3_2
INADL_3
RADIL
PTPN13_2
LNX2_2
MPDZ_3
DLG2_1
DLG2_2
PSMD9
NHERF1_2
DFNB31_3
PARD3B_1
MPDZ_7
PARD3_1
DLG1_2
TJP1_2
PDZD2_6
MAGI3_5
SHROOM2
SCRIB_4
GRIP2_1
GORASP1
GOPC
LNX1_2
PARD3B_3
SCRIB_3
MAGI1_2
MAST3
FRMPD3
MPDZ_13
TJP1_1
MAGI2_6
NHERF4_4
NHERF4_1
MAGI2_2
INADL_6
DLG1_1
SNTA1
NHERF3_4
DFNB31_1
MLLT4
HTRA4
SNTB1
PDZD7_1
SYNPO2
SNTG2
SNTG1
PARD3_2
PDLIM4
SCRIB_2
PDZRN3_1
SNTB2
TJP2_1
PTPN3
APBA1_2
MAGI3_3
DFNB31_2
FRMPD2_2
MYO18A
LIN7C
MAGI1_3
MAGI1_4
MAGI2_3
DLG2_3
DLG1_3
GRIP1_6
MPDZ_2
MPDZ_6
NHERF3_2
PPP1R9B
RIMS2
RIMS1
TJP3_3
MPP4
DLG4_3
DLG3_3
PTPN13_4
PDLIM2
MPP1
PDZRN4_1
DVL3
GRIP2_6
MAGI3_4
DLG3_1
TIAM2
ARHGAP21
PDZD2_3
PDZD2_1
LIMK2
USH1C_1
FRMPD2_1
TJP2_2
DLG5_3
DLG5_2
SYNPO2L
TJP3_1
LIN7B
GRIP2_7
IL16_1
MPP7
NOS1
PARD3B_2
MPDZ_10
PDLIM5
MPDZ_1
PDZD2_4
MAGI3_6
PDZD4
PDZD8
LNX2_3
PTPN4
LIN7A
LDB3
LAP2
INTU
INADL_4
USH1C_2
pkd
4.8
5.2
5.6
6.0
6.4
interactions
2
4
6
8
10
Aer_2
Aer_3
Aer_5
Bar_1
Bur_2
Bur_4
Ced_1
Chl_1
Chl_2
Cit_2
Cit_3
Cit_4
Ent_1
Esc_1
Esc_2
Esc_4
Esc_5
Esc_6
Esc_7
Esc_10
Esc_11
Esc_12
Esc_9
Kle_2
Leg_1
Leg_2
Mor_1
Phy_1
Pro_1
Pro_2
Pro_3
Pro_4
Pse_1
Pse_2
Sal_1
Sal_4
Ser_1
Shi_1
Shi_2
Shi_3
Shi_4
Shi_5
Vib_1
Vib_2
Vib_3
Yer_1
Yer_2
Yer_3
Yer_4
Yok_1
met_1
strain_abb
pkd_median
4.6
4.8
5.0
5.2
5.4
Distinct_interactions
1
15
30
45
60
0 5 10 15 20 25 30
Number of strains

**Supplementary Figure 8 | PDZ targeting by strain for interactions with pKd>4.5. a,** Interactions between individual PDZ domains and different bacterial strains grouped by genera on the x-axis (see supplementary table 1 for strain abbreviations) with vertical grid lines colored purple and orange for resident gut microbiota and known pathogen strains, respectively and **b,** total number of strains targeting each PDZ domain (above the specified affinity threshold). In both plots individual points are colored by median pkd and sized according to number of interactions with different PBMs. The y-axis is shared for a) and b).

#### ΔΔG against top human binders

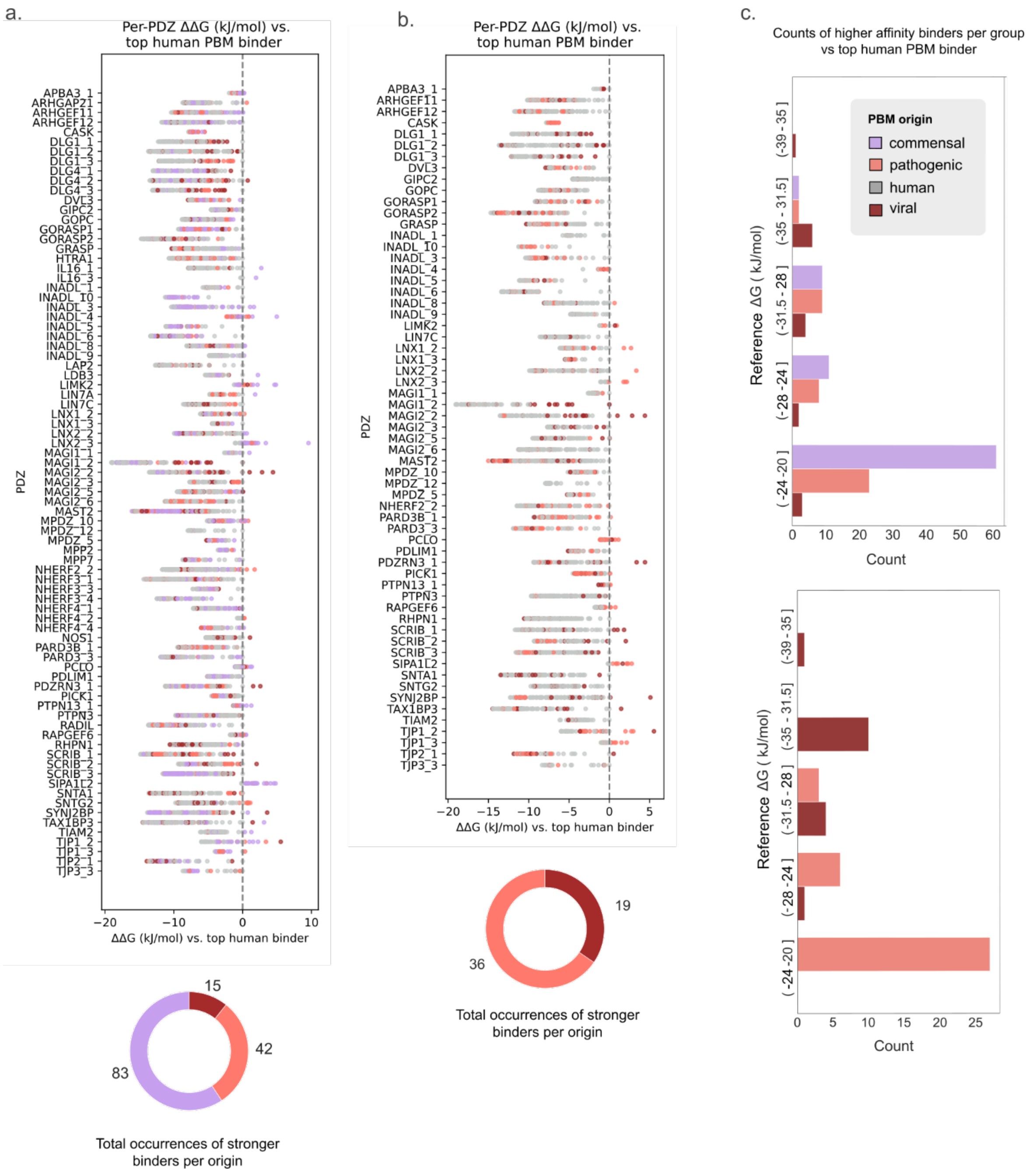


**Supplementary Figure 9 | Binding energy differences (ΔΔG) for PBMs of different origin against the top human PBM binder for each PDZ domain. a,** (top) scatter plot of the differences in binding energies (ΔΔG in kJ/mol) for PBMs from bacteria with gut tropism and human proteins expressed in gut tissues, with respect to the human PBM with the highest affinity for each PDZ domain. (bottom) Donut plot showing number of instances where a PBM of non-human origin has a higher affinity than the top human PBM binder for all PDZ domains combined **b,** Same as “a” but for lung tissues. **c,** Histograms showing counts of stronger binders of non-human origin with respect to the reference ΔG (in kJ/mol) for the highest affinity human binder for different reference ΔG intervals (stronger reference interactions on top of Y-axis) for gut (top) and lung (bottom) tissues.

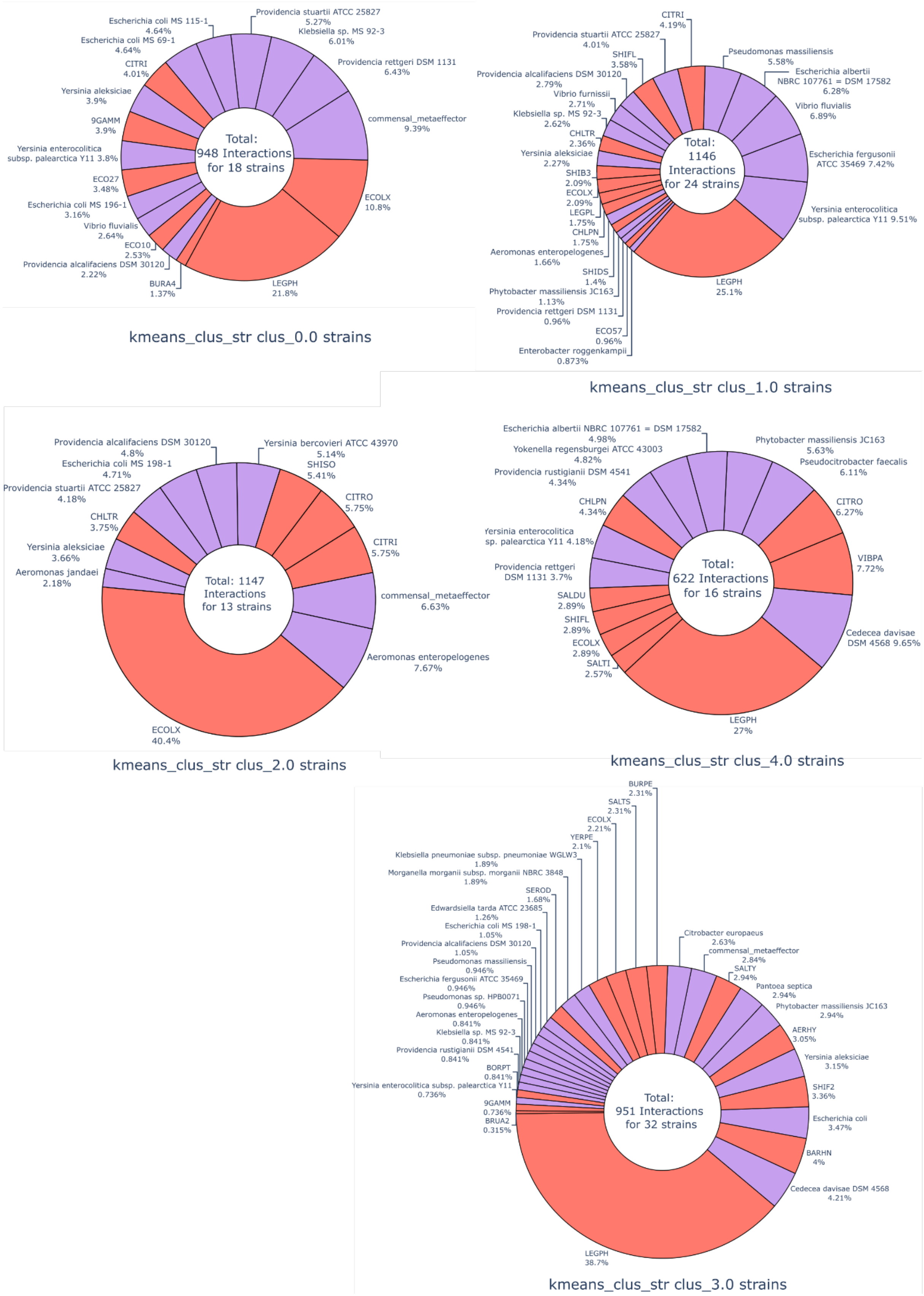


**Supplementary Figure 10 | Interaction similarity kmeans clusters strain composition**. Strain composition for each cluster from the interaction similarity analysis (Main Figure 2). Cluster numbers indicated at the bottom of each donut.

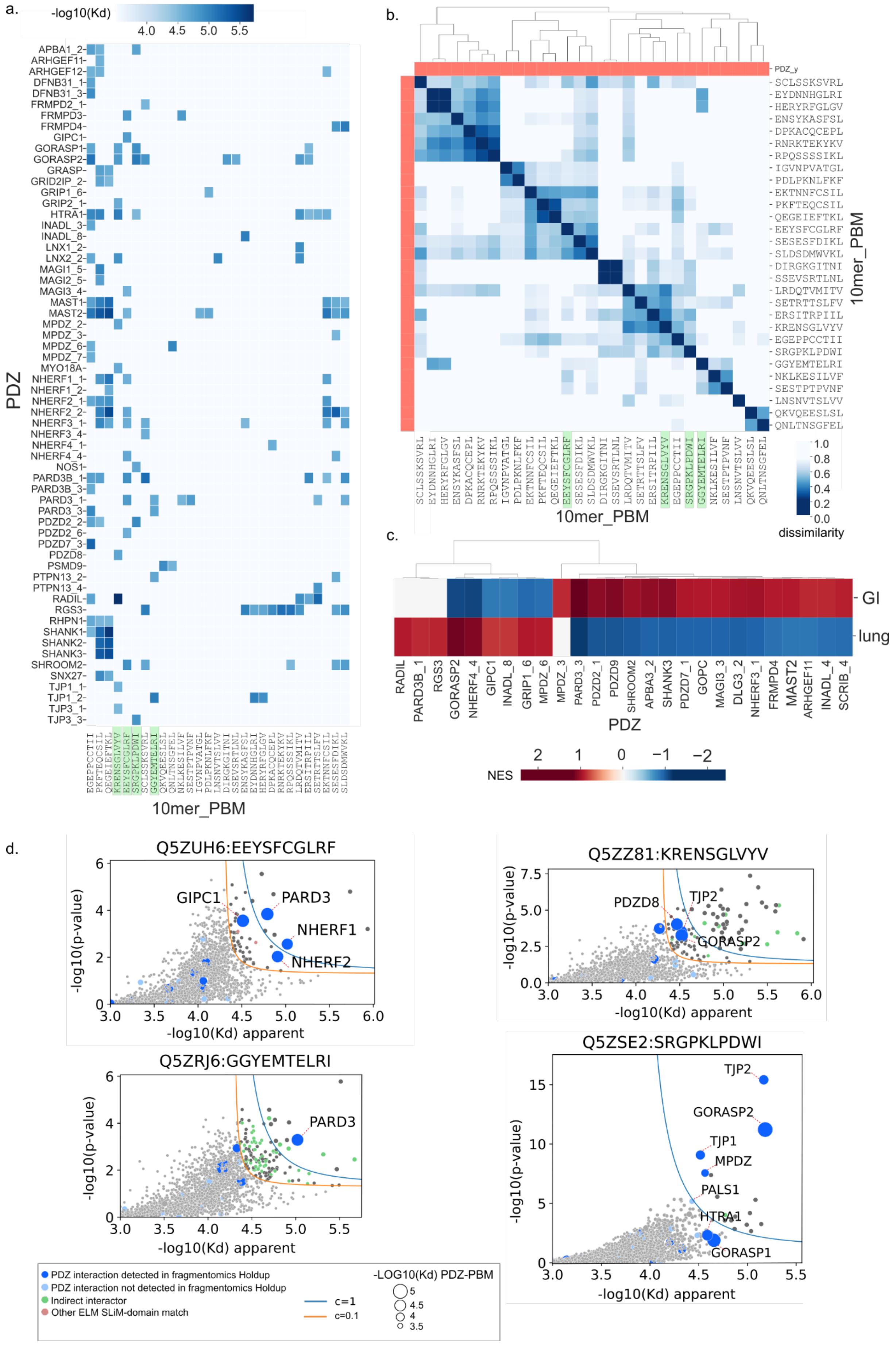

**Supplementary Figure 11 | *Legionella* PBM analysis (interactions with pKd >4.5).**
**a,** heatmap for all *Legionella* interactions with a pkd >4.5 and **b,** interaction dissimilarity

clustering (0 for highly similar profiles) for PBMs in a. **c,** GSEA normalized enrichment scores for preferential targeting of PDZ domains by tropism (gut and lung) **d.** native Holdup runs apparent -log10(Kd) scatter plots for four *Legionella* PBMs as baits against HEK293 cell extracts. MS for native Holdup for the bottom right PBM was performed with an Orbitrap Astral, and for the other three with an Orbitrap Exploris (see methods). Hyperbolic threshold curves calculated as in[1]. Navy blue, sky blue, green, salmon and grey colored dots represent PDZ proteins depleted by Holdup, PDZ proteins not detected by Holdup, indirect interactors of a depleted PDZ protein (annotated from IntAct[2] binary PPI data), proteins with Pfam[3] domains that recognize a motif other than the PBM on the bait peptide and depleted proteins not fitting into the previous groups, respectively.

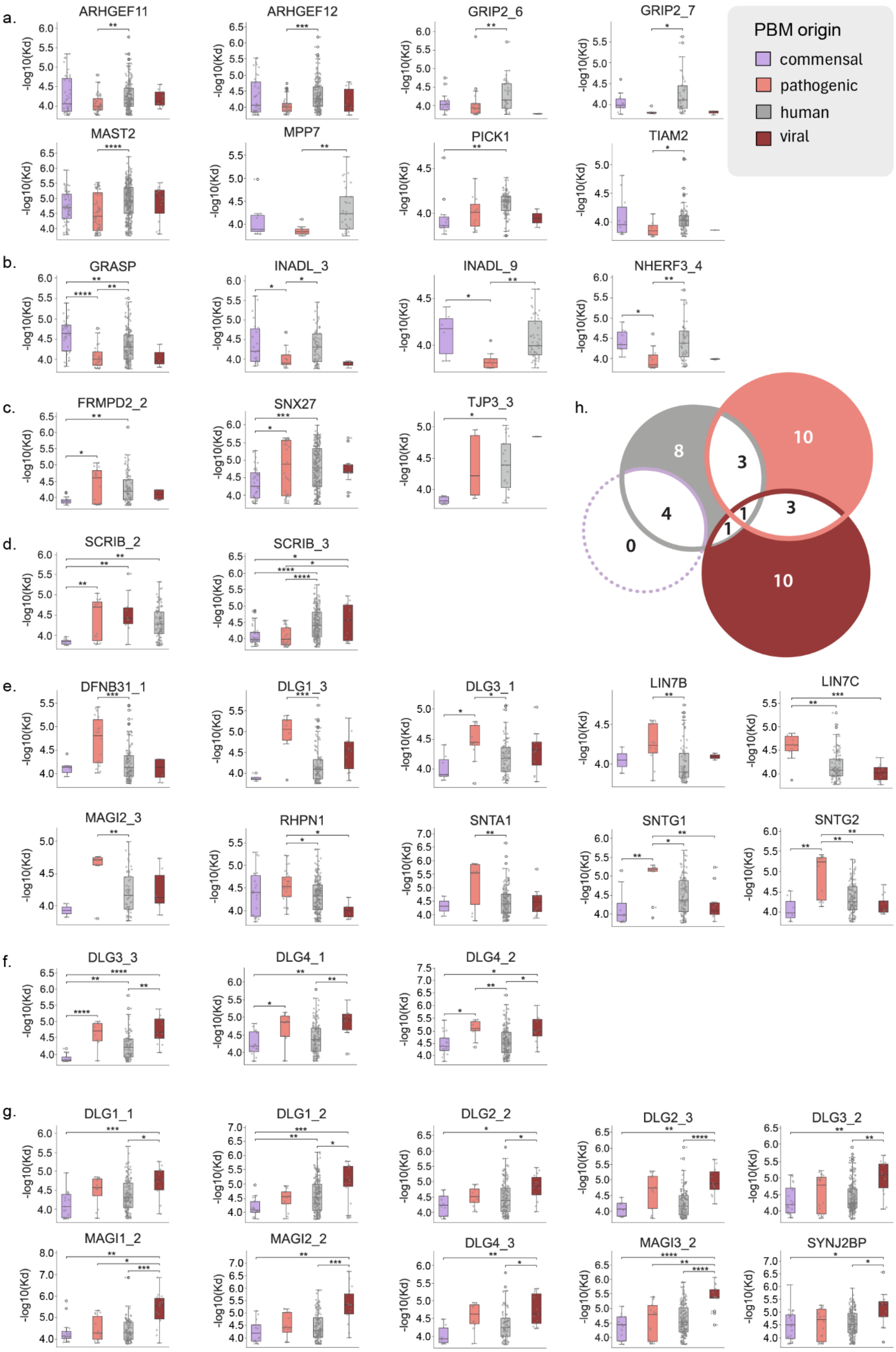


**Supplementary Figure. 12| Individual domain targeting by PDZ-binding motifs of different origins.** PDZ domains for which affinity distributions are significantly

(Kruskal-Wallis, Dunn’s correction, p-values<=0.05) higher for **a,** human PBMs exclusively compared to at least one other group **b,** human and commensal PBMs compared to bacterial pathogen PBMs, **c,** human and bacterial pathogen PBMs compared to commensal PBMs, **d,** (left) human, bacterial pathogen and viral PBMs compared to commensal PBMs and (right) human and viral PBMs compared to commensal and pathogen PBMs, **e,** bacterial pathogen PBMs exclusively compared to at least one other group, **f,** bacterial pathogen and viral PBMs compared to human and commensal PBMs (for DLG4_2), viral PBMs compared to human ones along with viral and pathogen PBMs compared to commensal ones (DLG4_1) and viral PBMs compared to human ones along with viral, human and pathogen PBMs compared to commensal ones (DLG4_3) and finally **g,** viral PBMs compared to at least one other group. **h,** Venn diagram showing the number of PDZ for which the distribution of affinities is significantly higher for PBMs of each origin compared to at least one other group along with overlaps for combined occurrences.

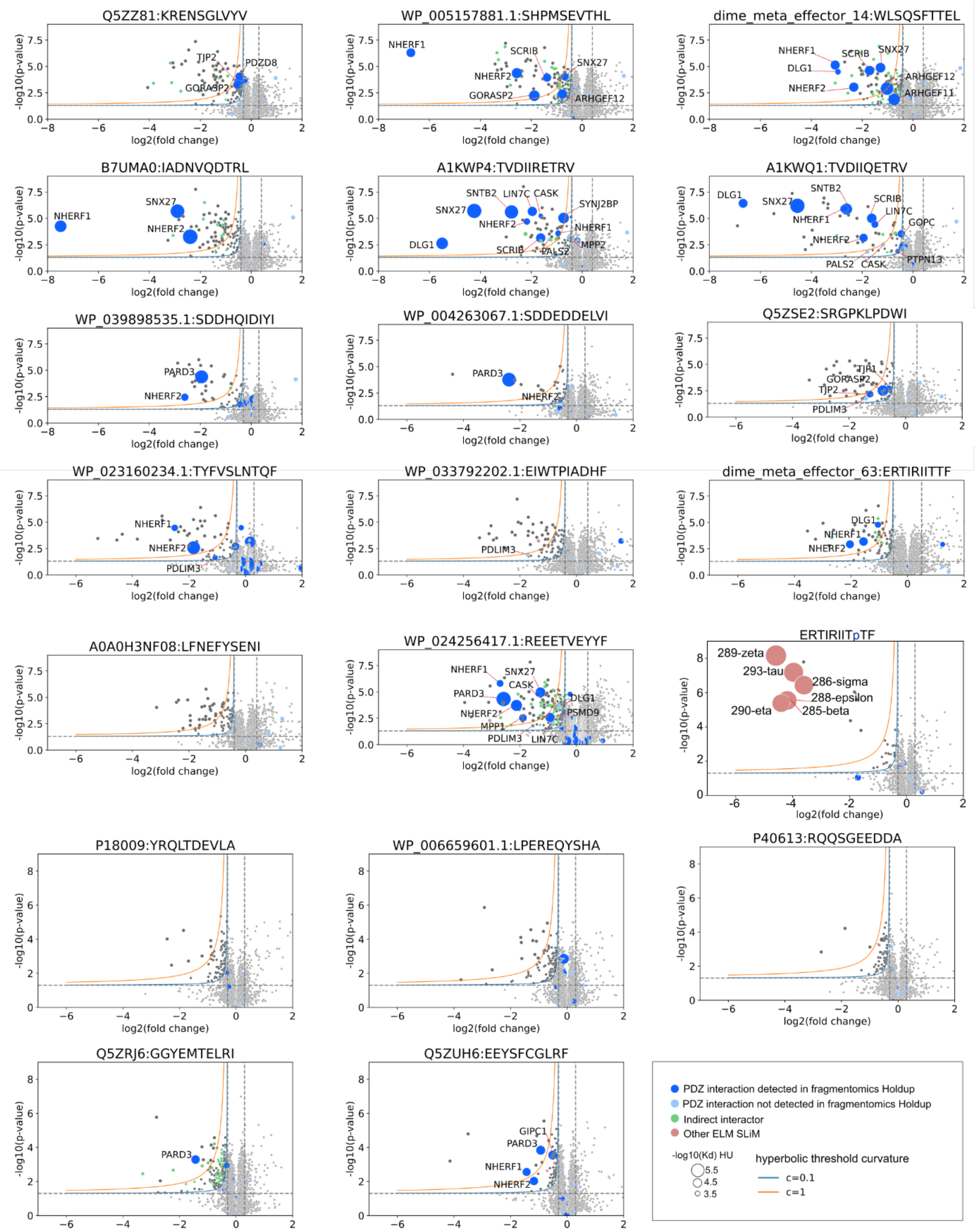


**Supplementary Figure 13 | Native Hold-up log2 fold change volcano plots for all groups.** Volcano plots of all (20) samples tested by Native Holdup with MS readout done by Orbitrap Exploris. The ERTIRIITpTF panel is the phosphorylated version (at position -1 from the C-terminal) of the panel above it (dime_meta_effector_63). Navy blue, sky blue, green, salmon and grey colored dots represent PDZ proteins depleted in fragmentomics Holdup, PDZ proteins not detected by fragmentomics Holdup, indirect interactors of a depleted PDZ protein (annotated from IntAct[2] binary PPI data), proteins with Pfam[3] domains that recognize a motif other than the PBM on the bait peptide and depleted proteins not fitting into the previous groups, respectively. Depleted PDZ protein gene names are annotated in each graph.

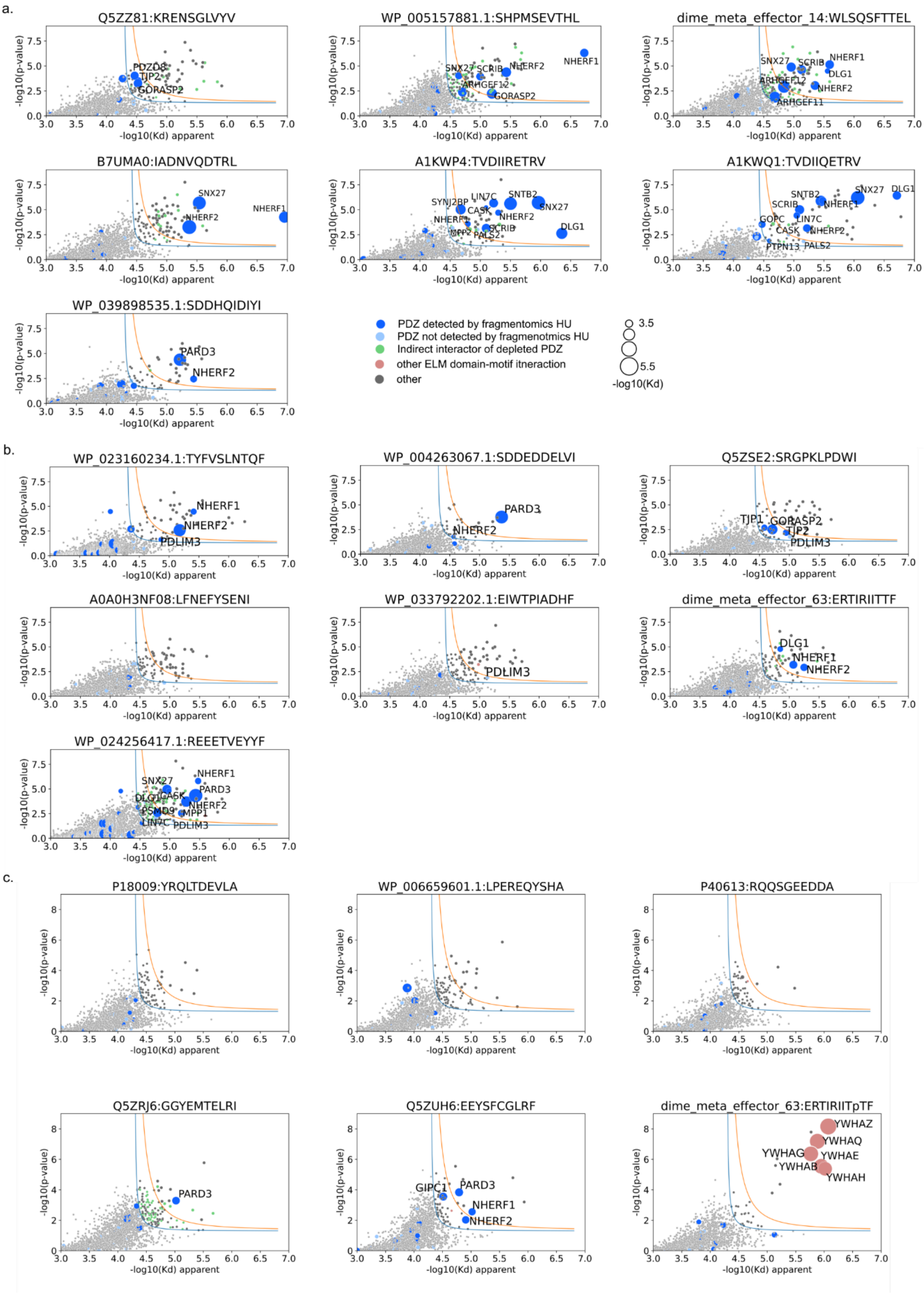


**Supplementary Figure 14 | Native Hold-up pkd scatter plots.** Scatter plots of apparent affinity for all 20 samples tested by native Holdup grouped by order of injection between negative controls. **a,** Group 1, **b,** Group 2, **c,** Group 3. Navy blue, sky blue, green, salmon and grey colored dots represent PDZ proteins depleted in fragmentomics Holdup, PDZ proteins not detected by fragmentomics Holdup, indirect interactors of a depleted PDZ protein (annotated from IntAct[2] binary PPI data), proteins with Pfam[3] domains that recognize a motif other than the PBM on the bait peptide and depleted proteins not fitting into the previous groups, respectively. Depleted PDZ and 14-3-3 protein gene names are annotated in each graph.

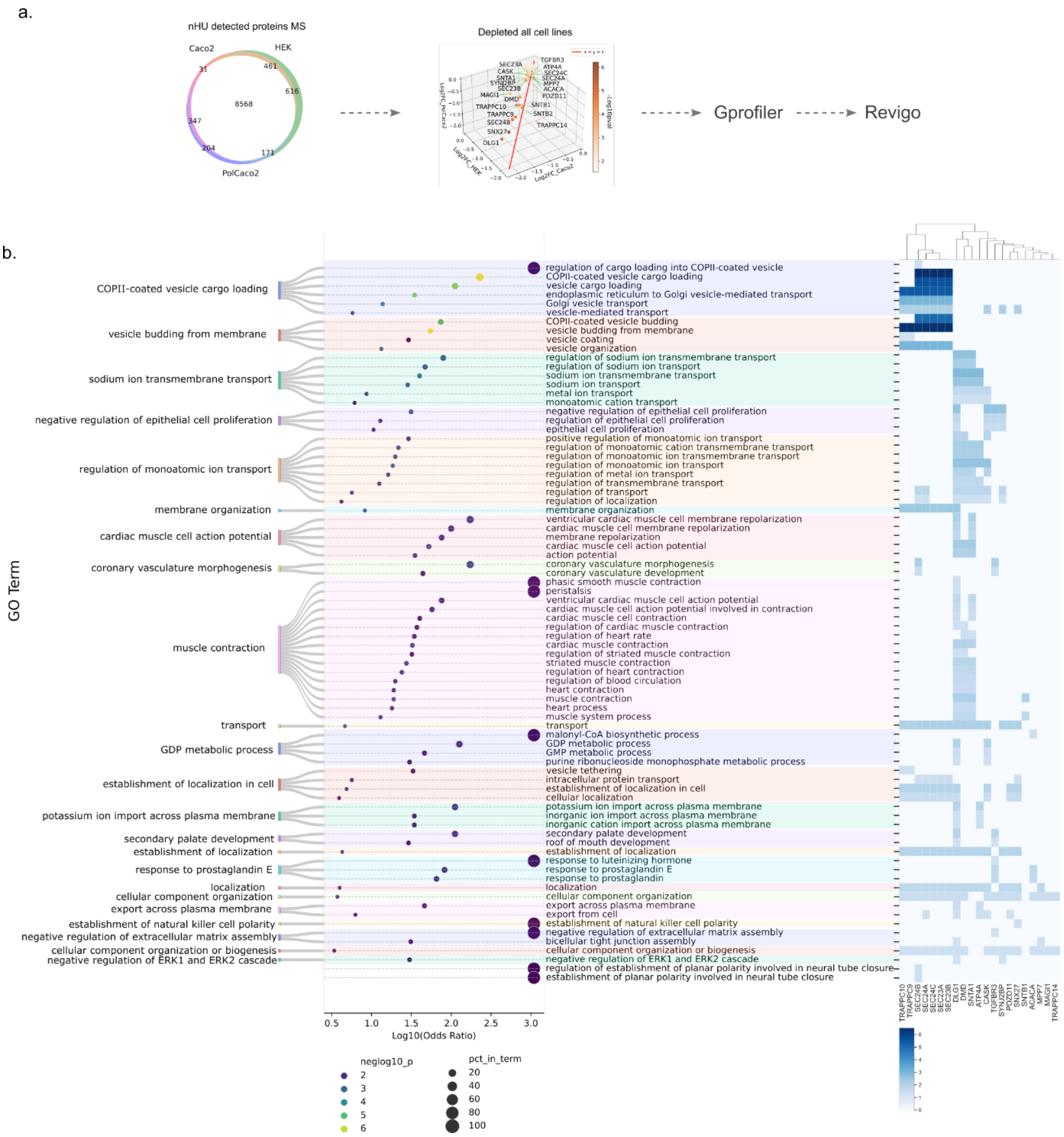


**Supplementary Figure 15 | Functional enrichment for full length NleA significant binders. a.** Functional enrichment workflow: (left) Venn diagram showing number of proteins detected by MS runs for each cell line/condition, with 8568 proteins in common in all cell lines. Scatter plot with proteins significantly depleted in all runs (log2(FC)<-0.2 and p-value<0.05) for full length NleA. Gprofiler enrichment analysis for depleted proteins in all runs was run against the background of proteins common to all runs (8568 in venn diagram) with p-value threshold at 0.01 and Benjamini-Hochberg correction. **b.** Revigo clusters (left) with associated log10 odds ratio bubble plot showing detailed log10 p value thresholds and percentage of query items in GO groups for individual GO terms.

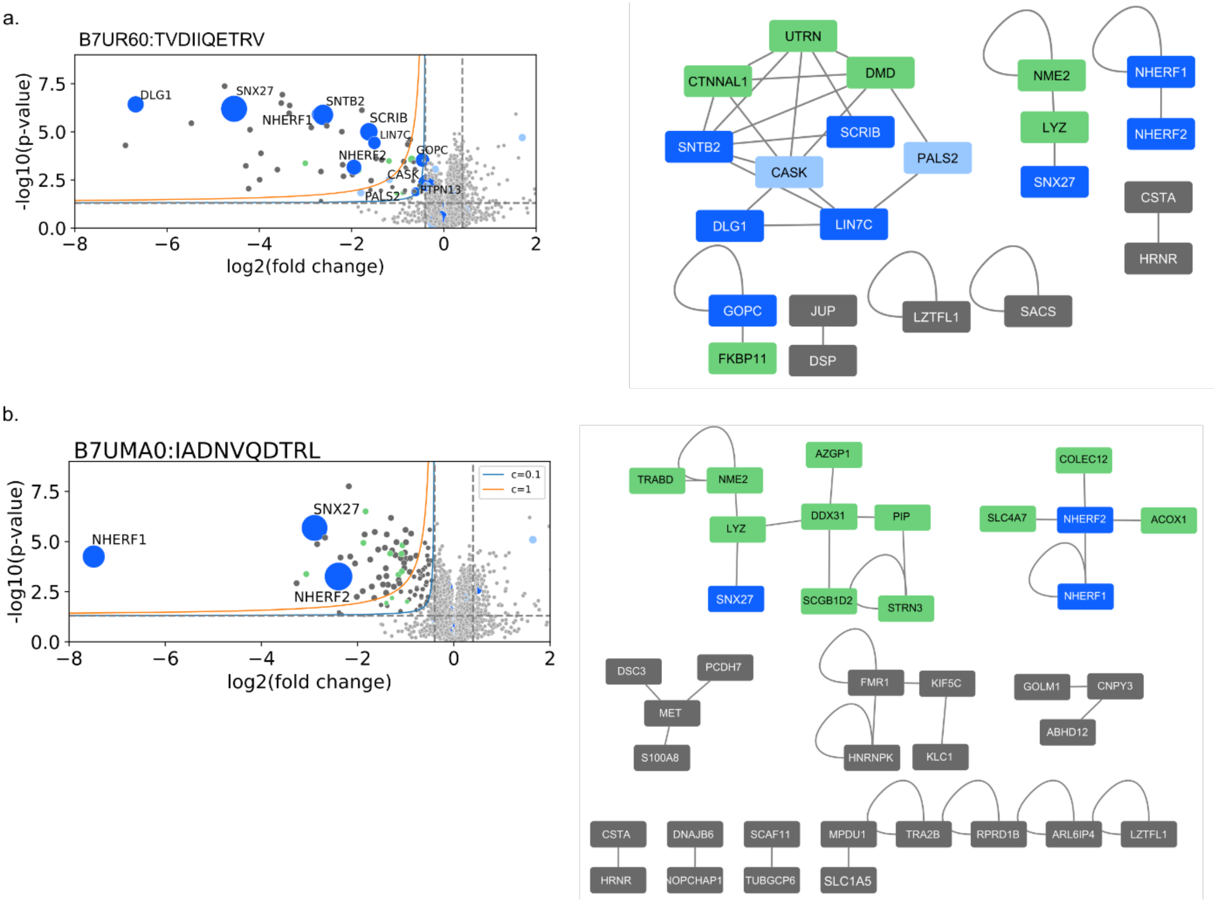


**Supplementary Figure 16 | Native Holdup and orthogonal binary PPI data from IntAct. a,** Native Holdup depletion profile for NleA PBM (left) along with interaction network of depleted protein region constructed from orthogonal data (IntAct[2]) of binary full length human PPIs. **b,** same as a) but for effector Map PBM.

#### Supplementary Tables

#### Supplementary Table 1 | Strain abbreviations

| Strain_id | Strain abbreviation Uniprot |
|---|---|
| Esc_1 | Escherichia albertii NBRC 107761 = DSM 17582 |
| Esc_2 | Escherichia fergusonii ATCC 35469 |
| Esc_3 | Escherichia coli |
| Esc_4 | Escherichia coli MS 115-1 |
| Esc_5 | Escherichia coli MS 198-1 |
| Esc_6 | Escherichia coli MS 69-1 |
| Esc_7 | Escherichia coli MS 196-1 |
| Esc_8 | Escherichia coli MS 21-1 |
| Esc_9 | ECOLX |
| Esc_10 | ECO27 |
| Esc_11 | ECO10 |
| Esc_12 | ECO57 |
| Pro_1 | Providencia alcalifaciens DSM 30120 |
| Pro_2 | Providencia stuartii ATCC 25827 |
| Pro_3 | Providencia rustigianii DSM 4541 |
| Pro_4 | Providencia rettgeri DSM 1131 |
| Yer_1 | Yersinia aleksiciae |
| Yer_2 | Yersinia enterocolitica subsp. palearctica Y11 |
| Yer_3 | Yersinia bercovieri ATCC 43970 |
| Yer_4 | YERPE |
| Vib_1 | Vibrio fluvialis |
| Vib_2 | Vibrio furnissii |
| Vib_3 | VIBPA |
| Aer_1 | Aeromonas dhakensis |
| Aer_2 | Aeromonas enteropelogenes |
| Aer_3 | Aeromonas jandaei |
| Aer_4 | Aeromonas tecta |
| Aer_5 | 9GAMM |
| Aer_6 | AERHY |
| Kle_1 | Klebsiella pneumoniae subsp. pneumoniae WGLW3 |
| Kle_2 | Klebsiella sp. MS 92-3 |
| met_1 | commensal_metaeffector |
| Pse_1 | Pseudomonas massiliensis |
| Pse_2 | Pseudomonas sp. HPB0071 |
| Pse_3 | Pseudomonas aeruginosa |
| Cit_1 | Citrobacter pasteurii |
| Cit_2 | Citrobacter europaeus |
| Cit_3 | CITRO |

| | |
|---|---|
| Cit_4 | CITRI |
| Phy_1 | Phytobacter massiliensis JC163 |
| Mor_1 | Morganella morganii subsp. morganii NBRC 3848 |
| Ced_1 | Cedecea davisae DSM 4568 |
| Pse_1 | Pseudocitrobacter faecalis |
| Edw_1 | Edwardsiella tarda ATCC 23685 |
| Edw_2 | Edwardsiella tarda ATCC 15947 = NBRC 105688 |
| Ent_1 | Enterobacter roggenkampii |
| Pan_1 | Pantoea septica |
| Yok_1 | Yokenella regensburgei ATCC 43003 |
| Leg_1 | LEGPH |
| Leg_2 | LEGPL |
| Sal_1 | SALTI |
| Sal_2 | SALTY |
| Sal_3 | SALTS |
| Sal_4 | SALDU |
| Shi_1 | SHIFL |
| Shi_2 | SHIF2 |
| Shi_3 | SHIB3 |
| Shi_4 | SHISO |
| Shi_5 | SHIDS |
| Bur_1 | BURTA |
| Bur_2 | BURPE |
| Bur_3 | BURP6 |
| Bur_4 | BURA4 |
| Chl_1 | CHLTR |
| Chl_2 | CHLPN |
| Clo_1 | 9CLOT |
| Bar_1 | BARHN |
| Bor_1 | BORPT |
| Cox_1 | COXBN |
| Ser_1 | SEROD |
| Och_1 | OCHA4 |
| Bru_1 | BRUA2 |

1. Gogl, G. *et al.* Quantitative fragmentomics allow affinity mapping of interactomes. *Nature Communications* **13**, 5472 (2022).
2. Del Toro, N. *et al.* The IntAct database: efficient access to fine-grained molecular interaction data. *Nucleic Acids Res* **50**, D648–D653 (2022).
3. Mistry, J. *et al.* Pfam: The protein families database in 2021. *Nucleic Acids Res* **49**, D412–D419 (2021).
